# Cell Type Specific Enhancers for Dorsolateral Prefrontal Cortex

**DOI:** 10.1101/2024.12.01.626253

**Authors:** Jing He, BaDoi N. Phan, Willa G. Kerkhoff, Aydin Alikaya, Olivia R. Brull, J. Megan Fredericks, Tao Hong, Morgan Sedorovitz, Chaitanya Srinivasan, Michael J. Leone, Olivia M. Wirfel, Samuel Dauby, Rachel K. Tittle, Meng K. Lin, Andreea C. Bostan, Bryan M. Hooks, Omar A. Gharbawie, Leah C. Byrne, Andreas R. Pfenning, William R. Stauffer

## Abstract

The dorsolateral prefrontal cortex (DLPFC) is crucial to primate cognitive functions, but a paucity of cell type specific tools limits studies of DLPFC neurocomputational principles. Therefore, we set out to identify enhancers that fit inside Adeno-Associated Virus (AAV) vectors and that elicited functional, cell type specific gene expression in the non-human primate (NHP) DLPFC. We used single nucleus RNA-Seq and ATAC-Seq from rhesus macaque tissue samples to define DLPFC cell types and their associated open chromatin regions (OCRs). We trained machine learning (ML) models to recognize the unique regulatory grammar associated with each DLPFC neuron type, performed in silico screening of all OCRs, and identified candidate enhancers most likely to elicit cell type specific transgene expression in each neuron type. For layer 3 pyramidal neurons (L3PNs) and layer 5 extratelencephalic neurons (L5ETs), we cloned the top twelve identified candidates into AAVs and injected them into NHP DLPFC. In situ observation of enhancer-driven expression revealed the best performers, RMacL3-01 and RMacL5ET-01. We validated RMacL3-01 and RMacL5ET-01 using one-at-a-time injections in NHP DLPFC. RMacL3-01 restricted GFP expression to pyramidal neurons in layers 2 and 3, whereas RMacL5ET-01 restricted expression to *POU3F1+* neurons in layer 5. RMacL3-01 elicited functional levels of channelrhodopsin expression that enabled optical activation of single- and multi-unit activity in NHP DLFPC. Together, these results and resources establish a solid foundation to study cell type specific principles of primate cognitive functions.

## INTRODUCTION

Human and nonhuman primates (NHPs) are capable of complex mental operations, including reasoning, inference, and sophisticated social cognition.^1–10^ The primate-specific dorsolateral prefrontal cortex (DLPFC) is implicated in these and other forms of cognitive control.^11^ Pharmacological studies of NHP DLPFC have shed some light on cellular contributions to DLPFC functions,^12–14^ but these methods lack the spatial and temporal capabilities of circuit-breaking tools such as optogenetics. Moreover, there are few cell type specific vectors capable of delivering circuit breaking transgenes to the NHP DLPFC. This lack of cell type specific tools prevents in-depth circuit-breaking studies to characterize the computations underlying complex cognition and the neural circuit dysfunctions underlying neuropsychiatric and age-related diseases, including schizophrenia and Alzheimer’s disease.

Cell type specific gene expression is controlled by transcription factors (TF) binding at proximal promoters and distal enhancers.^15–17^ Several NHP studies have exploited proximal promoters, including the tyrosine hydroxylase promoter or the L7 gene promoter, to drive the expression of channelrhodopsin (ChR2) in dopamine neurons and cerebellar Purkinje cells, respectively.^18–20^ Despite the promise of promoters demonstrated by these and other studies,^21^ systematic testing has revealed limited specificity in promoter-driven gene expression: promoters can drive broad, neuron class specific, but not neuron subtype specific, gene expression.^22^ In contrast, distal regulatory sequences, known as enhancers, show extraordinary promise to unlock circuits in wild-type animals, including NHP. Enhancers combine multiple TF binding sites compactly (∼500 bp) to achieve precise cell type specificity.^23^ When properly identified and packaged into Adeno-associated virus (AAV) vectors, enhancers can drive targeted gene expression in interneuron subtypes and layer specific neuron subtypes.^24–29^ Thus, enhancer-AAVs utilizing regulatory vocabulary matched to neuron subtype identity can be used to drive cell type specific transgene expression.

The major challenges are to properly identify cell type specific enhancers – the ENCODE database lists more than 1 million cell type specific and non-specific enhancers^30^ – and validate their cell type specific expression *in vivo*. Here, we collected snRNA-Seq and snATAC-Seq data from the DLPFC of adult rhesus monkeys to identify cell types and cell type specific open chromatin regions (OCRs). Using custom machine learning (ML) models, we identified and prioritized cell type specific OCRs as potential enhancers.^26,31^ The most promising enhancer candidates were synthesized and screened in low-complexity libraries for layer 3 pyramidal neurons (L3PNs) and Layer 5 extratelencephalic pyramidal neurons (L5ETs). The top performing enhancer, RMacL3-01, effectively restricted expression to layers 2 and 3. The top L5ET enhancer was specific for *POU3F1*+ L5ET neurons. RMacL3-01 successfully drove ChR2 expression *in vivo*; evoking functional neuronal activity in macaque DLPFC. These results lay the groundwork for investigating the neurophysiological contributions of DLPFC subtypes to NHP cognition.

## RESULTS

### DLPFC neuron phenotypes

To define DLPFC neuronal phenotypes that could be targeted by enhancers, we dissected fresh tissue from the DLPFC in five rhesus monkeys. The rostral and caudal extents of the anterior and posterior supra-principal dimples guided our dissections (Figure 1A, top). We cut coronal sections around these landmarks and separated the cortical gray matter from the underlying white matter (Figure 1A, bottom). We used well-validated single cell pipelines to perform single nucleus RNA with sequencing (snRNA-Seq) and single nucleus assay for transposase-accessible chromatin with sequencing (snATAC-Seq) (STAR Methods). We used the transcriptomic data from snRNA-Seq to define cell types and subtypes, and we used the open chromatin data from snATAC-Seq to identify enhancers to target those cell types and subtypes.

**Figure 1.**
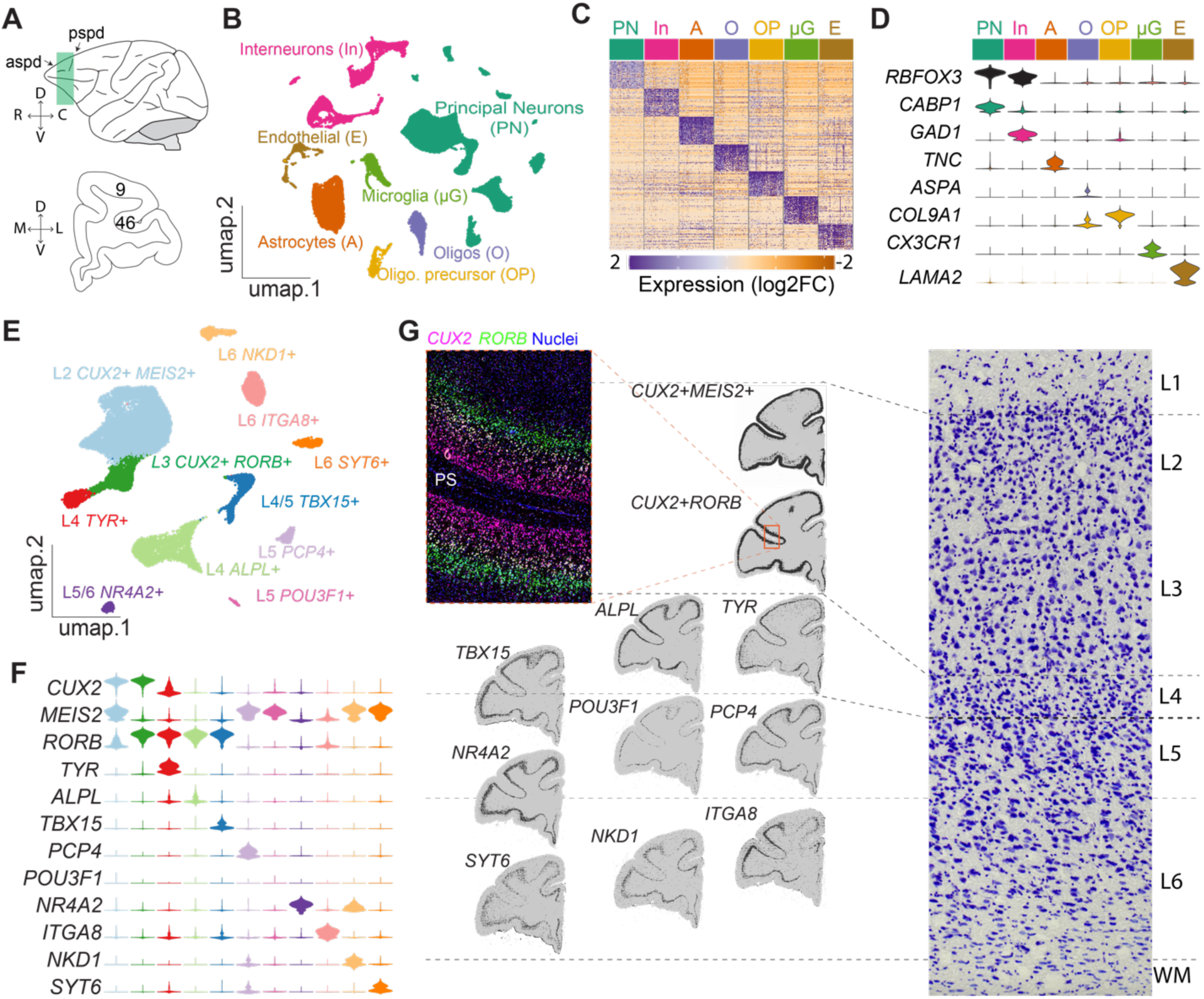
Rhesus macaque dorsolateral prefrontal cortex principal neuron types. (A) Dissected DLPFC regions. The green shaded area in between aspd and pspd indicates sample location. Axes show dorsal (D), ventral (V), rostral (R), caudal (C), medial (M), and lateral (L). (B) UMAP plot showing clustering of neuronal and non-neuronal DLPFC snRNA-Seq data from 5 rhesus monkeys. (C) Heatmap showing marker gene expression in each cell class. (D) Violin plot showing marker gene expression in each cell class. (E) Uniform Manifold Approximation & Projection (UMAP) plot showing 11 distinct excitatory subtypes in DLPFC. (F) Violin plot showing relative normalized marker gene expression in each excitatory cell type. Color match (E). (G) Fluorescence in situ hybridization (FISH) profiling of laminar organization of each excitatory cell type. The left enlarged image shows the original FISH signals in the red-boxed area. The right Nissl image indicates layer boundaries. *aspd*, anterior supra-principal dimple; *pspd* posterior supra-principal dimple; PS: principal sulcus. See also Figures S1 and S2.

Uniform Manifold Approximation and Projection (UMAP) dimensionality reduction revealed well-separated clusters (Figure 1B), and each of the five biological replicates contributed nuclei to each cluster (Figure S1A). Based on cluster-specific differential marker genes (Figure 1C), we identified seven major cell classes, including principal excitatory neurons (PN), inhibitory interneurons (In), astrocytes (A), microglia (μG), oligodendrocytes (O), oligodendrocyte precursors (OP), and endothelial cells (E). Violin plots revealed class specific marker gene expression levels (Figure 1D). Together, these results indicate that the dataset contains five high-quality biological replicates from the rhesus macaque DLPFC.

Our primary objective was to find enhancers that target the principal neurons of the DLPFC. We defined principal neuron clusters as those enriched for marker genes of excitatory neurons, including *SLC17A7* (Figure S1B).^32–34^ We re-analyzed this subset and identified 11 distinct excitatory subtype clusters (Figure 1E). We used differential gene analysis to identify excitatory subtype specific marker genes for each cluster (Figures 1F and S1C), and we used fluorescent in situ hybridization (FISH) against those marker genes to identify the anatomical position of each of those subtypes (Figure 1G). The FISH analysis revealed that the 11 clusters mapped onto subtypes that were layer specific and subtypes that spanned adjacent layers. Layer specific subtypes included L3PNs that expressed marker genes *CUX2* and *RORB* and L5PNs that expressed marker gene *POU3F1*. Subtypes that spanned adjacent layers included a layer 4/5 subtype that expressed *TBX15* and a layer 5/6 subtype that expressed *NR4A2* (Figure 1G). Hierarchical clustering and cosine similarity measures both showed that shallow- and deep-layer subtypes were molecularly separable, and that superficial layer subtypes were more similar to one another whereas layer 5 subtypes were highly distinct (Figure S1D) (p < 0.0001, Permutation tests, STAR Methods). These excitatory neuron subtypes closely corresponded to cell type taxonomies previously defined in humans, including near projecting (NP), cortico-thalamic (CT), inter-telencephalic (IT), and extra-telencephalic (ET) or previously described distinct cell types such as L6b and *Car3*+ L6IT PNs (Figures S1E-S1G).^35,36^ We will use these labels to define cell type specific OCRs.

We followed a similar approach to identify inhibitory interneuron subtypes in the rhesus macaque DLPFC. Analysis of inhibitory neuron clusters revealed 10 distinct subtypes (Figure S2A), which segregated based on their developmental origins from either the medial ganglionic eminence, marked by *LHX6* expression, or the caudal ganglionic eminence, marked by *ADARB2* expression (Figures S2B and S2C). Differential gene analysis identified subtype-specific markers for molecularly distinct populations of LAMP5+ and SST+ interneurons, as well as a discrete population of PVALB+ basket cells that co-expressed TH (Figure S2C). We used multiplexed FISH on coronal DLPFC sections to determine the laminar distribution of these interneuron subtypes (Figure S2D). The FISH analysis revealed that some interneurons subtypes showed layer preferences. For example, *NDNF*+ interneurons were restricted to L1, *VIP*+ interneurons were enriched in superficial layers, and the *PVALB+*/*TH*+ interneuron subtype populated L5-6 (Figure S2E). The remaining subtypes were distributed across multiple cortical layers. Together with the principal neuron taxonomies, these results establish a comprehensive taxonomy of DLPFC neuron types that can be used to identify subtype-specific enhancers.

### Enhancer Identification

Cell type specific OCRs may contain cell type specific enhancers. We collected single nucleus OCR data using a multi-omic assay that revealed both the transcriptome and OCRs in each cell, and thereby enabled direct annotation of OCRs with the cell type labels (STAR Methods). We used cell type labels from above (Figure 1) to identify reproducible, cell type specific, open chromatin profiles for all excitatory and inhibitory neuron subtypes (Figures 2A and S2F) (STAR Methods). This process revealed thousands of cell type specific OCRs for each neuron type (3,738 ± 1,048, mean ± standard error).

**Figure 2.**
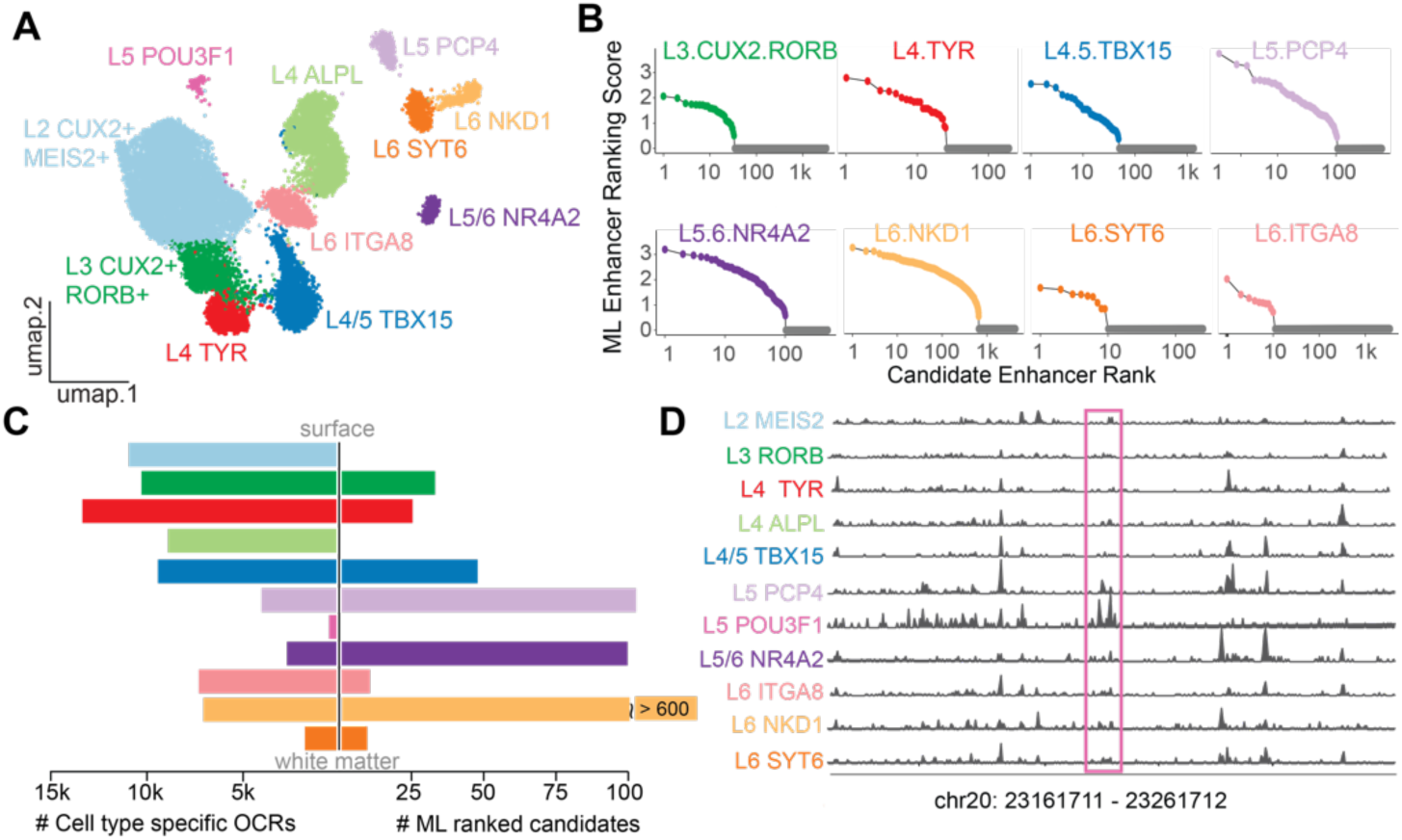
NHP DLPFC enhancer identification using single nuclear open chromatin (snATAC-seq) (A) UMAP plot showing clustering of excitatory neurons in DLPFC snATAC-Seq data from three rhesus monkeys. (B) ML ranking scores for OCRs in each cell type. (C) Cell type specific OCRs before ML ranking and selection (left plot), and after (right plot). Colors labeled in D. (D) Track plot showing an example peak for L5 *POU3F1*+ neurons. See also Figure S2.

Most OCRs are unlikely to elicit cell type specific gene expression. This is because most OCRs contain enhancers that lack the desired specificity, or that contain insufficient regulatory grammar to achieve desired levels of transgene expression.^29,37,38^ To efficiently screen for cell type specific enhancers, we previously developed an ML-based approach, SNAIL, to select OCRs for the capacity to elicit cell type specific gene expression in PV neurons relative to other neuron subtypes and glial cell types.^37^ Here, we generalized SNAIL to all neuron types. We used support vector machines (SVMs) and convolutional neural networks (CNNs) to train an ensemble of ML models for each neuron subtype (STAR Methods). Consistent with our original models for PV neurons,^37^ the areas under the receiver operator curves (auROC) ranged from 0.867 to 0.951 and the areas under the precision recall curves (auPRC) ranged from 0.876 to 0.958. We used the subtype ensembles to summarize the overall likelihoods for cell type specificity and off-target expression as ‘ML Enhancer Ranking Scores’ for all enhancer candidates (STAR Methods). We used the ML Enhancer Ranking Scores to prioritize candidates for *in vivo* testing (Figure 2B). The ML models substantially restricted the number of candidate enhancers for *in vivo* testing (3,738 ± 1,048 OCRs vs 233 ± 106 candidates; Figure 2C). However, using this stringent ML approach, we did not identify any candidate enhancers for the L5ET *POU3F1*+ neuron subtype. Therefore, we relaxed our criterion and paired it with domain heuristic approaches to select a panel of candidate enhancers for L5ET *POU3F1*+ neurons (Figure 2D, STAR Methods). The results of these ML-assisted rankings are testable sets of high-priority enhancer candidates.

### Enhancer screening

We selected the top twelve enhancer candidates for L3PNs and for L5ET *POU3F1*+ neurons for *in vivo* testing. We cloned the enhancers upstream of the HSP68 minimal promoter and placed GFP in the open reading frame (ORF). We included the HSP68 with GFP but no upstream enhancer as a control for the activity of the minimal promoter. We used human synapsin promoter (hSyn) with dTomato as a positive control. For each enhancer and the controls, we inserted a unique 500nt barcode downstream of the transgene. To ensure that all enhancers were at the same final titer, we packaged each plasmid individually into a backbone for AAV9-2YF. We then mixed the top 6 enhancer candidate AAVs, the positive control AAV, and the minimal promoter-only AAV, in equal fractions to create the L3PN screening library 1 (Figure 3A). We did the same procedure for the L3PN enhancers ranked 7-12, and for the L5ET enhancer candidates. This process resulted in two L3PN and two L5ET AAV screening libraries (STAR Methods, Table S1). The L3PN libraries were injected into the dorsal and ventral banks of the principal sulcus in one rhesus monkey (Figure 3B). The L5ET libraries were injected in the same locations, but in a different monkey. After one month, both animals were euthanized and DLPFCs were sectioned for FISH and immunohistochemistry.

**Figure 3.**
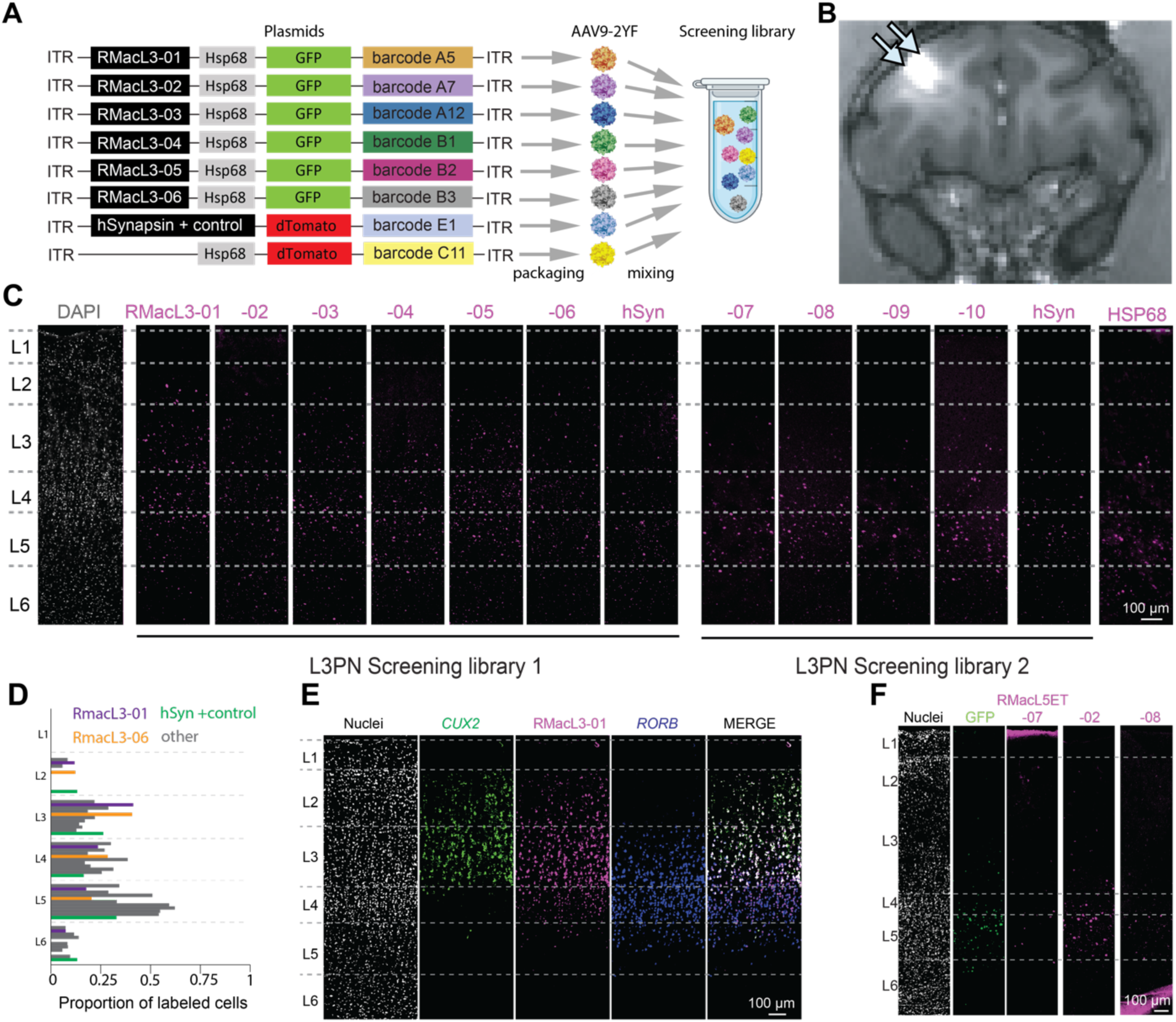
Screen of top enhancer candidates in NHP brain. (A) Cartoon summary of the screening procedure, L3PN screening library 1 shown as an example. We packaged each enhancer plasmid and the control plasmid individually into AAV9-2YF. We then mixed the top 6 enhancer candidate AAVs and the control AAV in equal fractions to create the L3PN screening library 1. (B) Post-injection MRI scanning of Gadolinium-doped AAV in the coronal plane clearly shows dorsal and ventral injection sites. (C) FISH analysis of ten enhancers and positive (hSynapsin promotor) and negative controls (minimal promotor HSP68 alone) driven transgene expression in monkey SM33. (D) Distribution of transgene expression in cortical layers. (E) Example FISH of RMacL3-01 driven expression with layer 2 and 3 excitatory neuron marker *CUX2* and layer 3 and layer 4 excitatory neuron marker *RORB*. (F) FISH analysis of L5ET enhancer candidates in monkey SM35. See also Figures S3 and S4.

We used FISH probes against the 500 nt, candidate-specific barcodes to measure enhancer performance (STAR Methods). On each section, we used 2-3 barcode probes, and we performed this on multiple adjacent sections to assess the layer-specificity of all candidate cell type enhancers and controls (Figure S3). Representative images of each enhancer candidate demonstrate that many were detected in superficial layers, but for many, especially in library 2, the detection level was low and centered on deep layers (Figure 3C). This deep-layer expression resembled the basal expression of the HSP68 minimal promoter when no enhancer was present (Figure 3C, right, Figure S3). For each AAV, we quantified the proportion of positive cells across DLPFC cortical layers by counting labeled nuclei in each layer (Figure 3D). This analysis revealed that, as expected, the hSyn positive control drove expression across layers 2-6, with a peak in layer 5. The analysis also highlighted two promising enhancer candidates, RMacL3-01 and −06. The most activity of both enhancers was centered on cells in layer 3. We selected RMacL3-01 – which was the sequence the ML models ranked first and had the strongest expression in L3 neurons– for deeper cell type profiling. In a subsequent round of FISH, we included marker genes for superficial (*CUX2*) and deeper (*RORB*) layers to identify layer 3 PN. This tissue processing showed that the majority of the RMacL3-01 signal was in L3PN, with some signal detected in both layers 2 and 4 (Figure 3E). Crosstalk between enhancer-AAVs has been documented,^39^ and given the subsequent results from one-at-time screening, this likely caused some of the non-specific labeling seen here. We repeated the FISH analysis for the L5ET enhancer candidates (Figures 3F, S4D, and S4E).

To further validate our screening approach, we investigated whether these primate-derived enhancers could maintain their layer specificity in rodents. Despite known differences in cortical organization between primates and rodents, particularly in upper layers, our pool of RMacL3 AAVs successfully drove layer 2/3-specific GFP expression in mouse cortex (Figures S4A-S4B), while the control hSyn promoter showed broad expression across cortical layers (Figure S4A). FISH analysis against enhancer-specific barcodes revealed that RMacL3-01 and RMacL3-02 were sufficient to drive layer 2/3-specific transcription in mouse cortex (Figure S4C). Similarly, the RMacL5ET AAV pool restricted GFP expression to mouse L5 (Figure S4D), with RMacL5ET-01 transcripts specifically co-localizing with the ET-specific marker *Fam84b* (Figure S4E). These promising results from both our multiplexed NHP screening and validation in mouse warranted rigorous testing of our top enhancer candidates, RMacL3-01 and RMacL5ET-01.

### One-at-a-time enhancer validation

We used one-at-a-time injections to validate the cell type specific expression of the best enhancers for L3PNs (RMacL3-01) and L5ETs (RMacL5ET-01). We packaged RMacL3-01 into three AAV9 variants, PHP.eB,^40^ 2YF,^41^ and X1.1.^42^ All injections were made directly into cortex via MRI-guided syringe pumps (STAR Methods, Table S2). In subject SM38, we injected the PHP.eB and 2YF vectors in the dorsal and ventral banks of the principal sulcus (Figures 4A and 4B). At high titer and close to the dorsal bank injection site, the PHP.eB vector resulted in non-specific GFP expression (Figure 4C). Further from the injection site, however, where the effective titer is lower, GFP expression was more specific to layers 2 and 3 (Figure 4C, inset). The 2YF vector generated highly restricted expression visible between the top of layer 2 and the granular layer 4 (Figures 4C-4E). Thus, enhancers in both AAV capsids were layer specific, but 2YF was more specific at the tested titer. In subject SM41, we injected the X1.1 and 2YF capsid variants into rostral and caudal aspects of the dorsal bank, respectively (Figure 4F). Inspection of the X1.1 injection site and surrounding tissue showed that the virus spread ∼3.5 mm in the rostro-caudal direction (Figure 4G). GFP expression appeared highly restricted to layers 2 and 3 (Figures 4G inset and 4J). We performed FISH against the barcode for RMacL3-01 and layer 4 marker *RORB*. We registered these images with the anatomical MRI and with the Nissl sections (Figure 4H). This enabled us to create a 3-D reconstruction of the expression derived from the X1.1 vector (Figure 4I). All these measures indicated that RMacL3-01 drove expression that was restricted to layers 2 and 3. In subject SM53, we injected three titers of the X1.1 vector (Figure 4K). We found that the highest titer resulted in non-specific expression, a tenfold dilution of that titer resulted in specific expression in layers 2 and 3, and a further 10-fold dilution was undetectable (Figures 4K-4M). In most cases, there was non-specific expression near visible injection sites, as previously observed.^26^ Together, these results indicate that enhancer RMacL3-01 - when injected at the appropriate titer (Figure 4L) - reproducibly restrict expression to layers 2 and 3.

**Figure 4.**
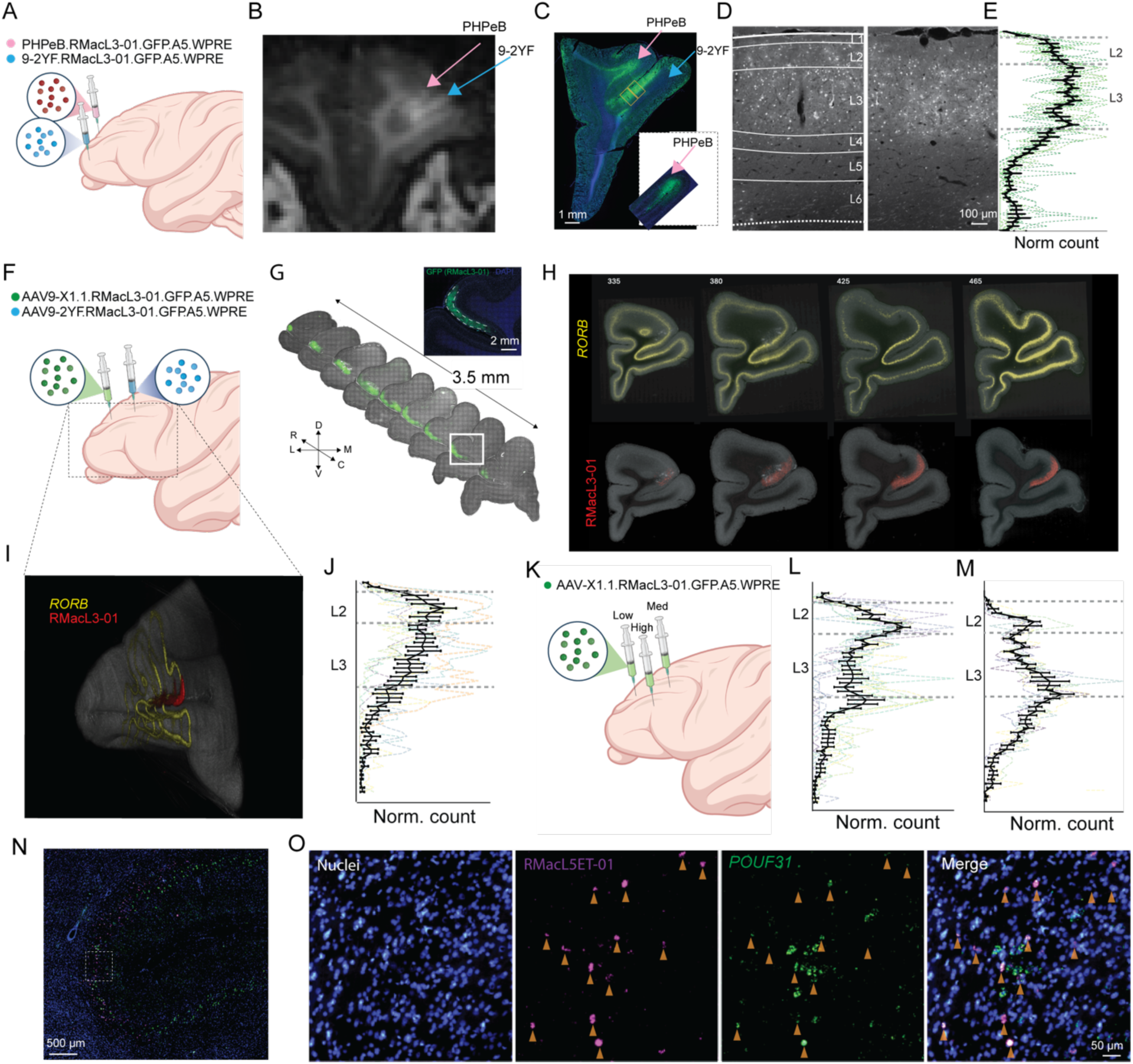
One-at-a-time validation of top enhancer candidates in NHP brain. (A) Cartoon showing injection sites in subject SM38 of RMacL3-01 packed in AAV9-2YF and PHP.eB. (B) Post-injection MRI scanning of Gadolinium shows the injection sites of RMacL3-01 packed in AAV9-2YF and PHP.eB. (C) Enhancer-driven GFP expression for RMacL3-01 packed in AAV9-2YF and PHP.eB. Inset shows layer-specific PHP.eB expression 1 mm caudal to the injection site. (D) High resolution images of enhancer-driven GFP expression, used for quantification, for RMacL3-01 packed in AAV9-2YF. (E) Layer distribution of enhancer-driven GFP expression for RMacL3-01 packed in AAV9-2YF. Dashed lines are counts of GFP+ cells from individual sections, and the black line is mean ± SEM. (F) Cartoon showing injection sites in subject SM41 of RMacL3-01 packed in AAV9 X1.1 and 2YF. (G) Rostral-caudal distribution of RMacL3-01-driven GFP expression. The inset is an enlarged image of the white box area. (H) FISH images aligned with the anatomical MRI and with the Nissl sections. (I) 3-D reconstruction of expression in subject SM41 in MRI space. (J) Layer distribution of enhancer-driven GFP expression for RMacL3-01 in SM41. Dashed lines are counts of individual sections, and the black line is mean ± SEM. (K) Cartoon showing SM53 injection of RMacL3-01 packed in AAV9 X1.1 at different titers. (L) Layer distribution of enhancer-driven GFP expression for RMacL3-01 at medium titer. Other conventions follow J. (M) Layer distribution of enhancer-driven GFP expression for RMacL3-01 at high titer. Other conventions follow J. (N) Dorsal and ventral banks of the cingulate sulcus with RMacL5ET-driven expression and L5ET marker *POU3F1*. (O) Enlarged view of (N) showing the colocalization of RMacL5ET-driven expression and L5ET marker *POU3F1*.

To quantify the cell type specificity of RMacL3-01, we used automated cell counting to examine the colocalization of the RMacL3-01 barcode with a known marker gene for layer 2/3 pyramidal neurons, *CUX2.* We did this on tissue sections from SM41 where we injected the X1.1 vector. The specificity for *CUX2* labeled cells was 77.2 ± 19.9% (mean ± SD) and the efficiency was 73.0 ± 12.4% (mean ± SD). This result indicates that not only was the enhancer layer specific, but it was also specific for layer 2/3 pyramidal neurons.

We packaged the most promising L5ET cell enhancer – RMacL5ET-01 – into AAV9-2YF and injected it into the ventral bank of the principal sulcus in subject SM39. As with RMacL3-01, we observed non-specific labeling near the injection site. However, farther from the injection site where the effective titer was diminished by diffusion, we observed highly restricted expression in L5 cells. In fact, we detected the highest specificity to layer 5 in medial wall structures around the cingulate sulcus, which is ∼3 mm from the injection site (Figure 4N). Multi-channel FISH against the RMacL5ET-01 barcode and a L5ET marker gene, *POUF31*, showed high colocalization (Figure 4O). This result indicates that RMacL5ET−01 can effectively target layer 5 projection neurons.

### Functional testing of RMacL3-01 with optogenetics

The enhancer-driven AAVs were developed to enable circuit-breaking studies in NHPs, especially with light-gated ion channels such as ChR2. Therefore, we performed *in vivo* validation using RMacL3-01 to drive expression of ChR2. We exchanged the GFP with ChR2(H134R)-p2A-GFP. The opsin-containing plasmids were packaged into AAV9-2YF and AAV9-X1.1 capsids and injected into the dorsal and ventral banks of the principal sulcus in SM63 (Figure 5A). Two months after injection, we opened a 2×3 cm cranial window to expose the principal and arcuate sulci (Figure 5A, STAR Methods). We used a handheld royal blue light, 500 nm long pass filter, and USB camera to visualize the injection sites (Figure 5B). We used a custom-made 16-channel linear electrode array with one lightguide and four windows; one window was located between electrodes 3 and 4, another between 6 and 7, 9 and 10, and 12 and 13. The lightguide was attached to a 473nm, 500 mW laser (STAR Methods). The array was approximately aligned to the principal sulcus (STAR Methods), aimed near injection sites, and advanced until spikes were detected on the most superficial channel (Ch 1). Optical pulse trains readily evoked activity along the length of the array (Figure 5C). The activity included multi-unit and single unit action potentials (Figure 5D). Optically evoked activity followed the frequency of the laser stimulus (Figure 5E). These results demonstrate that the RMacL3-01 enhancer can drive functional levels of ChR2 expression. Thus, RMacL3-01 vectors are suitable for immediate application to circuit-breaking studies in the NHP DLPFC.

**Figure 5.**
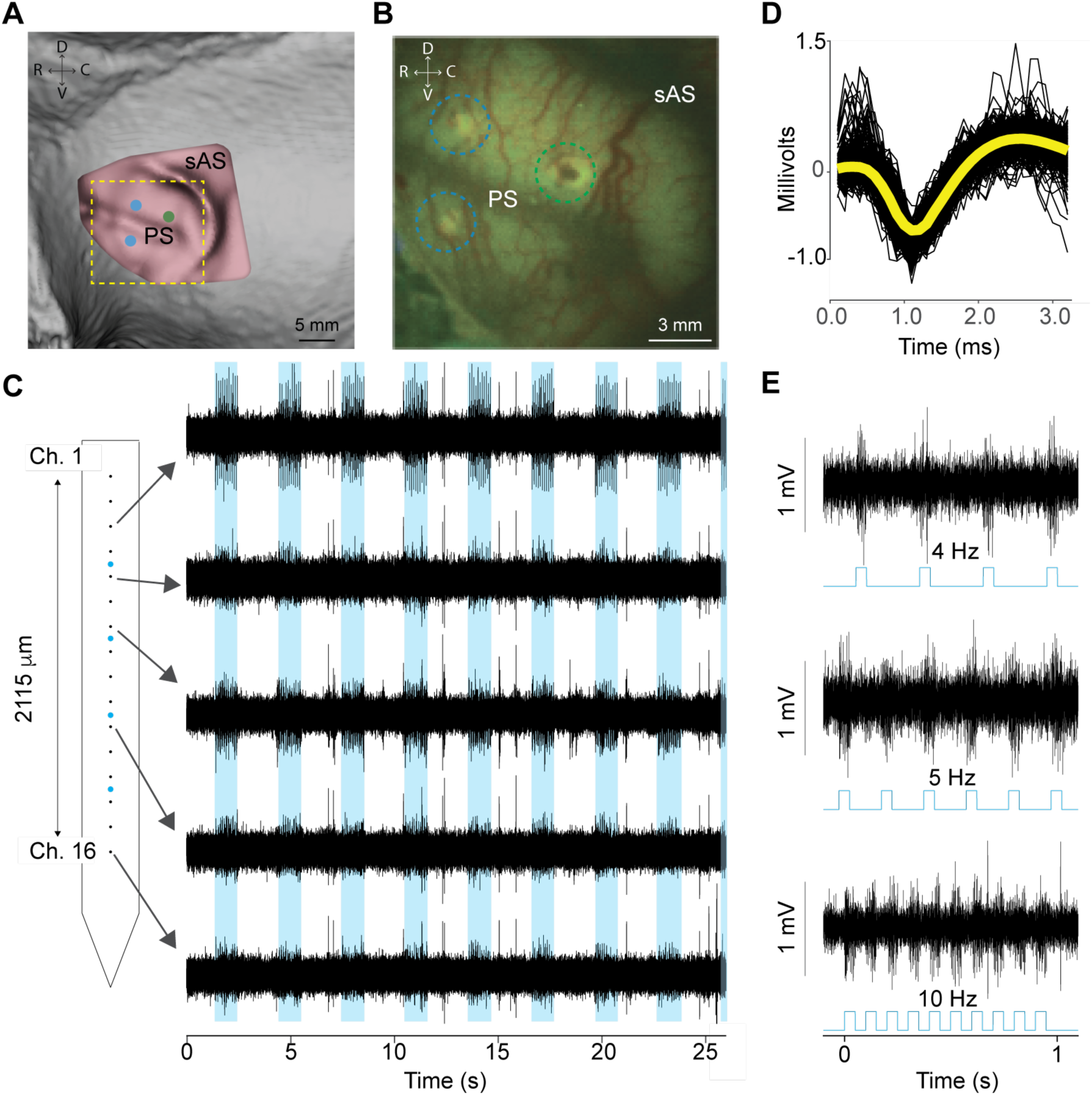
Optogenetic control of enhancer-driven ChR2 in NHP brain. (A) Cartoon modification of MRI scan, which depicts the 2×3 cm cranial window opened to expose the principal sulcus (PS) and arcuate sulcus (sAS), two months after injection. Injection sites are shown for RMacL3-01 driving ChR2, packaged into AAV9-2YF (rostral, blue dots) and AAV9-X1.1 (caudal, green dot). (B) Injection sites visualized in cortex with a handheld UV light and USB camera during the optical stimulation experiment. (C) Custom-made 16-channel linear electrode array with four windows. Simultaneous voltage traces recorded on channels 3, 5, 7, 11, and 16. Optical pulse trains (shown in blue) evoked neural activity along the length of the array. (D) Voltage traces of optically evoked single unit waveforms. The yellow curve is the average waveform. (E) Optically evoked activity followed the frequency of the laser stimulus, which was applied at 4, 5, and 10 Hz in the example traces.

## DISCUSSION

NHPs, and especially Old-World monkeys like rhesus macaques, present special challenges that limit circuit-based investigation of their cognitive functions. They are large and expensive. They reach sexual maturity between 3 (F) and 4 (M) years old, and females can generate at most one offspring per year. These factors seriously complicate the germline modifications and selective breeding technologies used to generate transgenic rodents. Nevertheless, rhesus macaques are irreplaceable as a model organism if we want to understand human cognition and disease.^43,44^ The present work was undertaken to overcome some of these challenges for a primate-specific brain region – the six-layered DLPFC.^45,46^ We used snRNA-Seq to uncover, identify, and define the molecular characteristics of DLPFC neuron subtypes in rhesus macaque monkeys (Figure 1). We applied these molecular phenotype definitions to snATAC-Seq data from rhesus macaque DLPFC and created custom-made ML models to discover and prioritize subtype specific enhancers (Figure 2). We cloned these enhancers into AAV9-based libraries together with enhancer-specific barcodes. We injected the libraries into rhesus and cynomolgus macaque brains and screened for layer and cell type specific gene expression (Figure 3). This screening revealed several candidate enhancers for layer 3 and 5 neurons (Figure 3). We performed one-at-a-time validation in multiple capsids and revealed that RMacL3-01 reliably targeted layer 2/3, albeit in a titer-dependent fashion. RMacL5ET-01 was highly specific to *POU3F1+* cells, also in a titer-dependent fashion (Figure 4). Finally, we demonstrated that RMacL3-01 drove functional levels of ChR2 (Figure 5). Together, our results provide a comprehensive resource that includes multi-omic single cell data from rhesus macaque DLPFC, several effective enhancers, and validated protocols for circuit-breaking studies of higher cognitive functions.

The NHP DLPFC has no direct homolog in rodents.^45–47^ Furthermore, there is substantial heterogeneity in cell type specific gene expression, even between different cortical regions within the same primate species.^33,48,49^ For these reasons, we collected single cell transcriptomic and epigenomic data from rhesus macaque DLPFC. We used transcriptomic data from five animals to define consensus cell types, and we identified reproducible OCRs across three animals for all excitatory and inhibitory neuron subtypes (Figures 2C and S2F). This dataset is a valuable resource for this and future studies. Although several other studies have collected transcriptomic data from DLPFC,^50^ NHP studies have small numbers of subjects. Thus, our additional biological replicates make an important contribution to the field by expanding the foundation of high-quality single-cell genomic data from rhesus macaques.

Cell type specific enhancers can be found within cell type specific OCRs. However, the number of cell type specific OCRs was high, ranging from thousands to tens of thousands, in the cell types of interest (Figure 2). This large number of OCRs creates an enormous search space, which renders exhaustive, *in vivo* screening for enhancers intractable. Moreover, most OCRs lack the genome sequence features needed to drive cell type specific expression.^26,29,38^ For these reasons, we created cutting-edge ML models that learned to recognize the genome sequence features that predicted cell type specific activity and ranked the OCRs according to their potential as cell type specific enhancers.^26,31^ Here, we scaled the ML models to screen for enhancers in all distinct neuronal subtypes in the NHP DLPFC. For all neuron subtypes with sufficient number of samples, model auROC and auPRC ranged from 0.867 to 0.958. Thus, these ML models are sufficiently sensitive and specific to screen hundreds of thousands of OCRs to nominate a few cell type specific regulatory DNA sequences that drive cell type specific transgene expression. In addition to the ML rankings – we used differential TFBS binding, proximity of open chromatin to marker gene, and inferred correlation of open chromatin to nearby transcription to refine enhancer predictions. These additional features made use of the multi-omic data collected here and further reduced the number of candidate enhancers for screening. These *in silico* screens resulted in a testable number of high-priority enhancer candidates for each cell type. Moreover, we note that RMacL3-01 was the top-ranked layer 3 enhancer. Thus, the enhancer rankings predicted success in eliciting cell type specific gene expression.

*In silico* screening produced a list of highly likely enhancer candidates, and we selected the most promising candidates for L3PNs and L5ET cells for *in vivo* screening. We created low-complexity libraries – libraries that included less than ten candidate enhancers – to screen for cell type specificity. Crucially, each enhancer candidate was paired with a DNA barcode that was detectable via FISH. This enabled us to screen all the selected enhancer candidates in a small number of monkeys. The results of screening the L3PN library did not reveal any enhancer candidates whose activities were completely restricted to layer 3. One potential cause of this lack of screening specificity was transcriptional crosstalk, a phenomenon where AAV episomes interact with one another inside the cell.^39^ AAV transcriptional crosstalk is a well-documented phenomenon^39,51^ and a major roadblock to large-scale enhancer screening. Nevertheless, the results we obtained in the screen suggested a few enhancers that should be validated with one-at-a-time injections, where enhancer crosstalk is not a factor. In one-at-a-time enhancer validation, we saw much higher layer specificity (Figure 4). In all cases, the top-ranked ‘layer 3’ enhancer showed expression across layers 2 and 3. This is likely because layer 2 and layer 3 pyramidal neurons are very similar from a transcriptomic perspective, a fact which can be appreciated by how closely adjacent they are in the cell type UMAP plot (Figure 1E). Additionally, in all cases, we found a very strong modulation of cell type specificity with titer and capsid. Despite this dependence on capsid and titer, we were able to reproducibly target L2/3 and L5 ET cells.

A further challenge to developing cell type specific, enhancer-driven AAVs is determining whether highly specific variants can be used to drive sufficient levels of circuit-breaking transgenes. This challenge is exacerbated in NHPs, where such testing often involves chair-training animals, implanting chronic or acute recording chambers, followed by months or years of neurophysiological testing. To accelerate this process, we pursued functional validation in terminal procedures. We injected enhancer-driven AAVs into naïve animals in the surgical theatre. After 1-2 months’ time for expression, the animal is brought back into the surgical theater and the brain is exposed under general anesthesia. This procedure enabled us to visualize the GFP expression, and to target optoelectrodes into sites with the highest levels of expression (Figure 5B). In doing so, we quickly verified that RMacL3-01 drove functional expression of ChR2. These results indicate that RMacL3-01 can be an effective tool for photo-identification of L3PN and for stimulation-based studies.

The current study has several limitations. One of the most important limitations was our selection of the HSP68 minimal promoter. Post-mortem analysis indicated that this promoter drove significant expression even in the absence of any enhancer (Figure 3C). Further studies that use alternative minimal promoters with less basal activity could reveal greater specificity and might, in fact, reveal other enhancers that are potent cell type specific drivers. Second, the enhancers were paired with the same DNA barcodes for all experiments in this study. These DNA barcodes were selected because the ML model predicted they did not possess enhancer activity. Nevertheless, we cannot be certain that the same pattern of enhancer-driven gene expression would occur in the absence of the barcode. To avoid this issue in future testing, we will use two or three barcodes per enhancer. Another limitation was that we were unable to control for crosstalk between co-injected constructs. Therefore, we have not completely explored the small space of the potential enhancers revealed by our ML models. Further method development to minimize crosstalk is a requirement for effective, high-throughput screening. Finally, our functional validation studies were done with one circuit-breaking transgene: ChR2. The application of other circuit-breaking tools, including inhibitory opsins, DREADDs, and genetically coded calcium sensors, will likely require more testing and development. Despite these limitations, we have identified, validated, and established effective protocols for enhancer-AAVs that drive cell type specific expression in L2/3 and L5ET neurons in NHP DLPFC. Further, we have established an effective pipeline using Rhesus macaque multi-omics data to develop more cell type and circuit specific AAVs.

Our ultimate goal is to understand the circuit specific computational principles that underlie sophisticated cognitive functions such as reasoning, deduction, and choice.^1,4–6^ To accomplish these goals, we require cell type specific AAVs that can deliver the experimental control offered by modern circuit breaking tools like optogenetics, chemogenetics, and optical activity sensors. Moreover, next generation circuit targeted gene therapy approaches will require viral vectors that work in large, wild-type primates. The present work is aimed at achieving both goals. Our high throughput *in silico* screen successfully prioritized effective enhancers with high hit rate, specificity, and expression, and greatly reduced experimental costs for validation. Our approach to functional validation in terminal procedures also provides a roadmap to functionally validate enhancers across the NHP brain. The outcomes include enhancers for cell types that are crucial for cognition and especially affected by aging and neuropsychiatric diseases.^50^ Moreover, the protocols and procedures used here provide a roadmap to building a substantial arsenal of cell type specific tools with which to understand high-level cognitive functions and deliver circuit-targeted therapies for neuropsychiatric and neurodegenerative disorders.

## DATA AVAILABILITY STATEMENT

The 24 enhancer-AAV constructs screened in this study (12 for L3PNs and 12 for L5ETs) and the construct pAAV.RMacL3-01.ChR2(H134R)-GFP will be available from Addgene at the time of publication. Datasets will be available at NCBI GEO at the time of publication. The codes to perform snATAC analysis and develop SNAIL ML models for this paper will be accessible from https://github.com/pfenninglab/Macaque_Multiome_DLPFC at the time of publication.

## ACKNOWLEDGEMENTS

The authors thank J. Breter for animal care and enrichment, N. Towler for histology, and A. Brown for molecular biology. This study utilized a Brainsight Turnkey Neuro-Navigation System for NHP Research, which was provided by National Institute of Mental Health grant S10MH126994 to W.R.S. This paper was supported by the NIH BRAIN Initiative Armamentarium for Precision Brain Cell Access through the following grants: National Institute of Mental Health UG3/UH3MH120094 to A.R.P, L.C.B, W.R.S, and UF1MH130881 to A.R.P, L.C.B, W.R.S. BNP was supported by National Institute on Drug Abuse grant F30DA053020.

## AUTHOR CONTRIBUTIONS

Conceptualization, L.C.B., A.R.P., and W.R.S.; Methodology, J.H., B.N.P., A.R.P., and W.R.S.; Software, J.H., B.N.P., C.S., M.J.L., and A.R.P.; Formal Analysis, J.H., B.N.P.,C.S., M.J.L., A.R.P., and W.R.S.; Investigation, J.H., B.N.P., W.G.K., A.A., O.R.B., J.M.F., T.H., M.S., O.M.W., S.D., M.K.L, and A.C.B., O.A.G., L.C.B., A.R.P., and W.R.S, Writing – Original Draft, J.H., B.N.P., A.R.P., and W.R.S.; Writing – Review & Editing, J.H., B.N.P., R.K.T., A.R.P., O.A.G., and W.R.S.; Funding Acquisition, L.C.T.B., A.R.P., and W.R.S.; Resources, B.M.H., A.R.P., and W.R.S.; Supervision, A.R.P. and W.R.S.

## DIVERSITY AND INCLUSION

One or more of the authors of this paper self-identifies as a member of the LGBTQ+ community.

## SUPPLEMENTAL TABLES

**Table S1.**
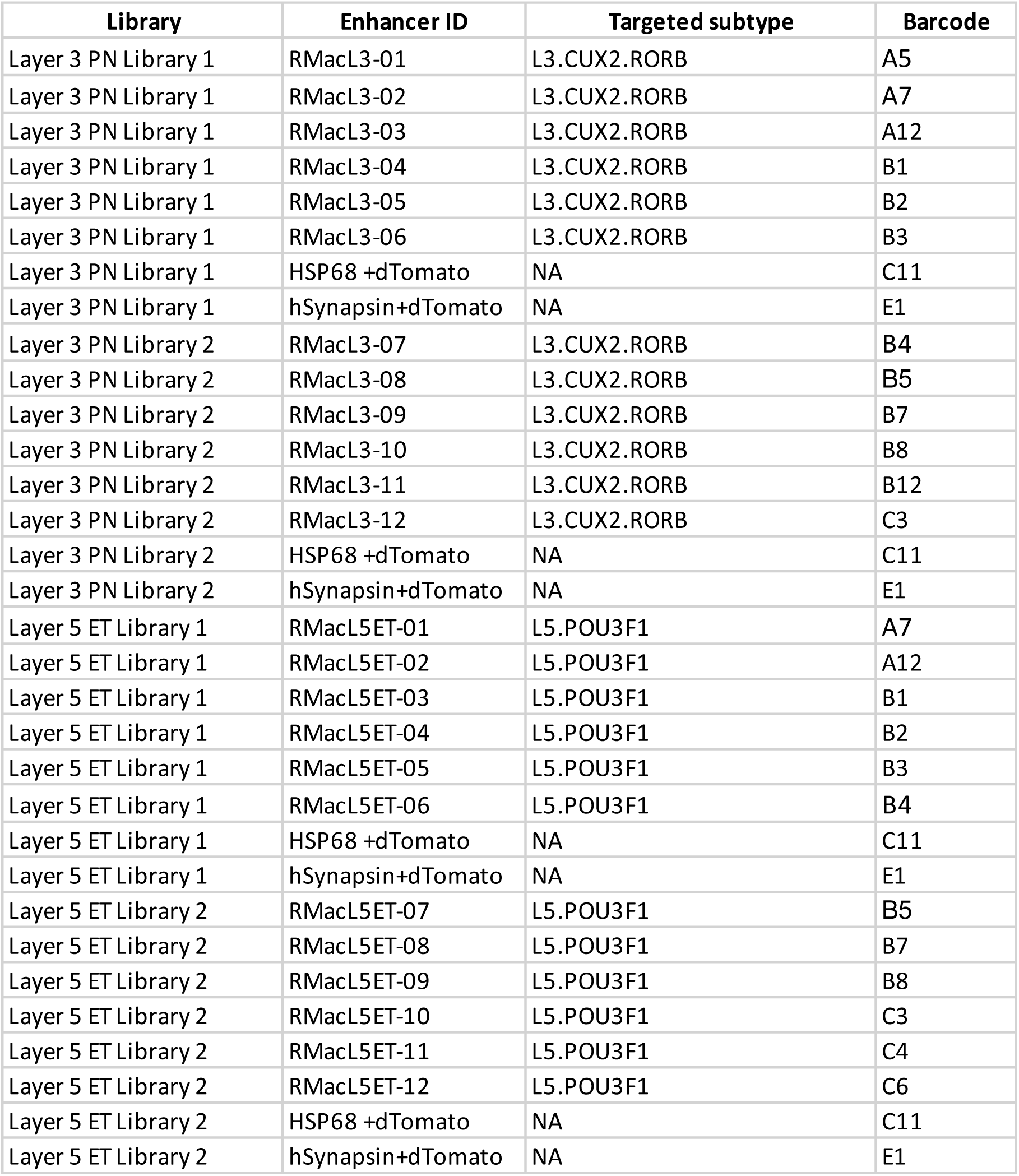
Summary of enhancer libraries mentioned in this paper, Related to Figures 3-5.

| Library | Enhancer ID | Targeted subtype | Barcode |
| --- | --- | --- | --- |
| Layer 3 PN Library 1 | RMacL3-01 | L3.CUX2.RORB | A5 |
| Layer 3 PN Library 1 | RMacL3-02 | L3.CUX2.RORB | A7 |
| Layer 3 PN Library 1 | RMacL3-03 | L3.CUX2.RORB | A12 |
| Layer 3 PN Library 1 | RMacL3-04 | L3.CUX2.RORB | B1 |
| Layer 3 PN Library 1 | RMacL3-05 | L3.CUX2.RORB | B2 |
| Layer 3 PN Library 1 | RMacL3-06 | L3.CUX2.RORB | B3 |
| Layer 3 PN Library 1 | HSP68 +dTomato | NA | C11 |
| Layer 3 PN Library 1 | hSynapsin+dTomato | NA | E1 |
| Layer 3 PN Library 2 | RMacL3-07 | L3.CUX2.RORB | B4 |
| Layer 3 PN Library 2 | RMacL3-08 | L3.CUX2.RORB | B5 |
| Layer 3 PN Library 2 | RMacL3-09 | L3.CUX2.RORB | B7 |
| Layer 3 PN Library 2 | RMacL3-10 | L3.CUX2.RORB | B8 |
| Layer 3 PN Library 2 | RMacL3-11 | L3.CUX2.RORB | B12 |
| Layer 3 PN Library 2 | RMacL3-12 | L3.CUX2.RORB | C3 |
| Layer 3 PN Library 2 | HSP68 +dTomato | NA | C11 |
| Layer 3 PN Library 2 | hSynapsin+dTomato | NA | E1 |
| Layer 5 ET Library 1 | RMacL5ET-01 | L5.POU3F1 | A7 |
| Layer 5 ET Library 1 | RMacL5ET-02 | L5.POU3F1 | A12 |
| Layer 5 ET Library 1 | RMacL5ET-03 | L5.POU3F1 | B1 |
| Layer 5 ET Library 1 | RMacL5ET-04 | L5.POU3F1 | B2 |
| Layer 5 ET Library 1 | RMacL5ET-05 | L5.POU3F1 | B3 |
| Layer 5 ET Library 1 | RMacL5ET-06 | L5.POU3F1 | B4 |
| Layer 5 ET Library 1 | HSP68 +dTomato | NA | C11 |
| Layer 5 ET Library 1 | hSynapsin+dTomato | NA | E1 |
| Layer 5 ET Library 2 | RMacL5ET-07 | L5.POU3F1 | B5 |
| Layer 5 ET Library 2 | RMacL5ET-08 | L5.POU3F1 | B7 |
| Layer 5 ET Library 2 | RMacL5ET-09 | L5.POU3F1 | B8 |
| Layer 5 ET Library 2 | RMacL5ET-10 | L5.POU3F1 | C3 |
| Layer 5 ET Library 2 | RMacL5ET-11 | L5.POU3F1 | C4 |
| Layer 5 ET Library 2 | RMacL5ET-12 | L5.POU3F1 | C6 |
| Layer 5 ET Library 2 | HSP68 +dTomato | NA | C11 |
| Layer 5 ET Library 2 | hSynapsin+dTomato | NA | E1 |

**Table S2.** Summary of enhancer library injections in monkeys mentioned in this paper, Related to Figures 3-4.

| **Subject** | **Enhancer** | **Serotype** | **Titer** | **Volume** |
| --- | --- | --- | --- | --- |
| SM38 | RMacL3-01 | PHP.eB | 6.7 E13 | 15 µl |
| SM38 | RMacL3-01 | 2YF | 4.5 E13 | 15 µl |
| SM41 | RMacL3-01 | X1.1 | 5.5 E13 | 15 µl |
| SM41 | RMacL3-01 | 2YF | 5.0 E13 | 15 µl |
| SM53 | RMacL3-01 | X1.1 | 1 E13 | 15 µl |
| SM53 | RMacL3-01 | X1.1 | 1 E12 | 15 µl |
| SM53 | RMacL3-01 | X1.1 | 1 E11 | 15 µl |
| SM39 | RMacL5ET-01 | 2YF | 5 E12 | 15 µl |

## SUPPLEMENTAL FIGURES

**Figure S1.**
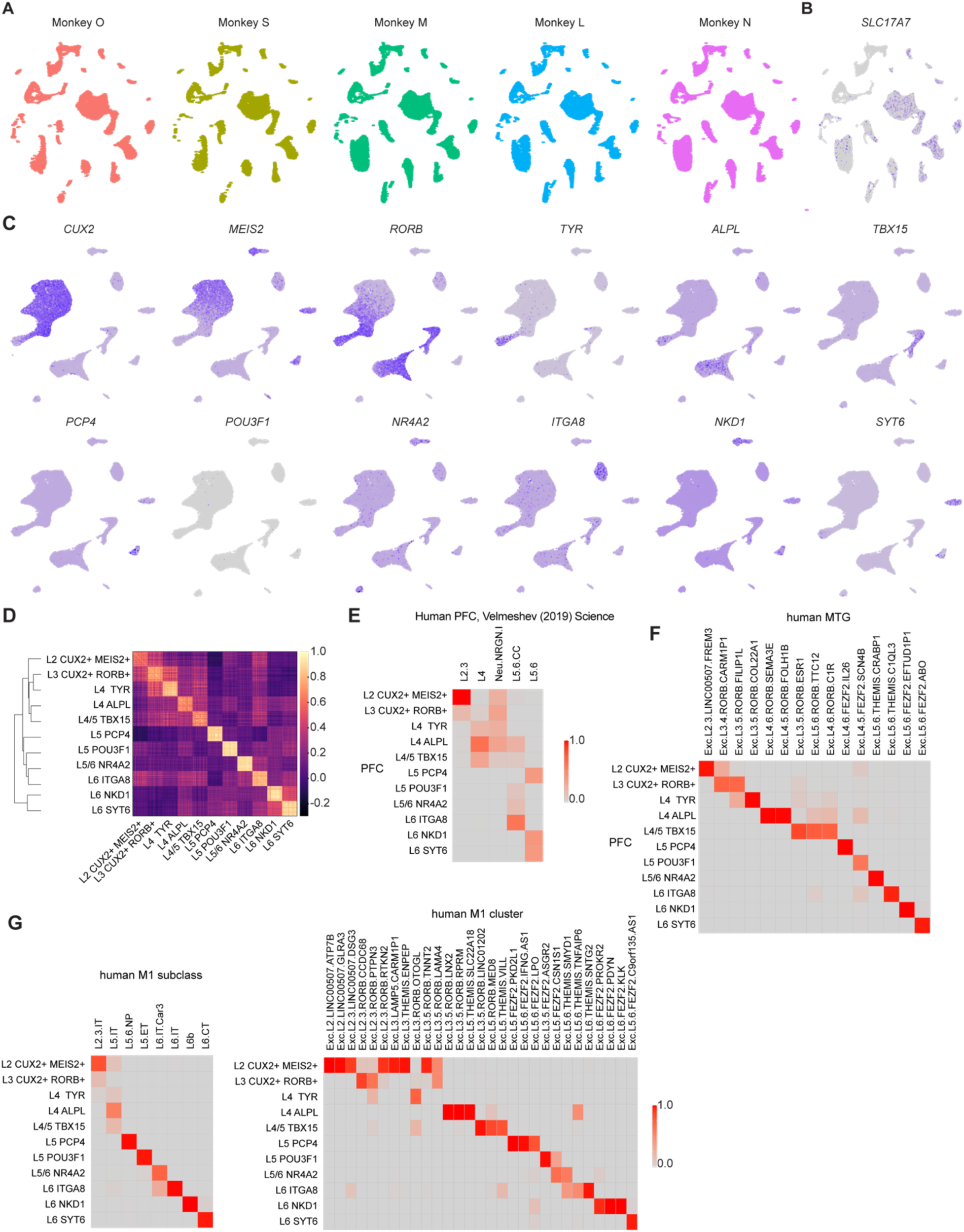
Excitatory neurons in primate cortex, Related to Figure 1. (A) UMAP plot showing clustering of neuronal and non-neuronal DLPFC snRNA-Seq data in 5 rhesus monkeys. (B) Feature plot of neuron-specific marker *SLC17A7*. (C) Feature plots of excitatory neuron subtype markers. (D) Heat map showing the cosine similarity within and between the eleven types of excitatory neurons. (E) Comparison of monkey excitatory neuron subtypes with human prefrontal cortex (PFC) dataset. (F) Comparison of monkey excitatory neuron subtypes with human medial temporal gyrus (MTG) dataset. (G) Comparison of monkey excitatory neuron subtypes with human primary motor cortex (M1) dataset.

**Figure S2.**
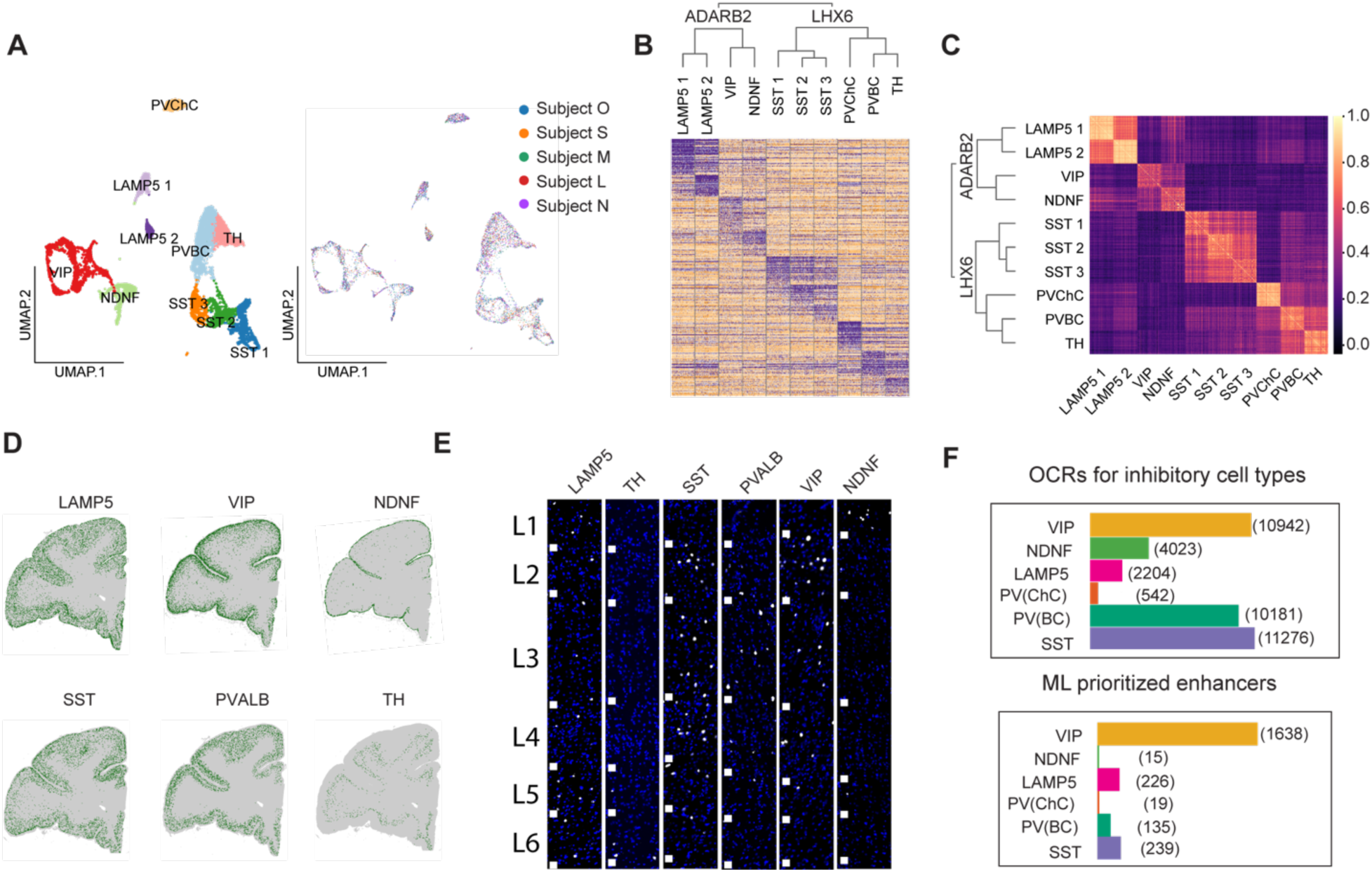
Inhibitory neurons in primate DLPFC, Related to Figures 1 and 2. (A) UMAP visualization of NHP DLPFC inhibitory neuron subtypes, colored according to subtypes (left) or individual subjects (right). (B) Heat map of differentially expressed genes. (C) Heat map showing the cosine similarity within and between the ten subtypes of inhibitory neurons. (D) Cell profiler processed marker gene expression in the PFC coronal sections. (E) FISH analysis of 6 major inhibitory cell types. White squares indicate cortical layer borders. (F) Cell type specific OCRs before and after ML ranking and selection for inhibitory neurons. The top plot shows the number of OCRs before ML ranking, and the bottom plot shows the number after.

**Figure S3.**
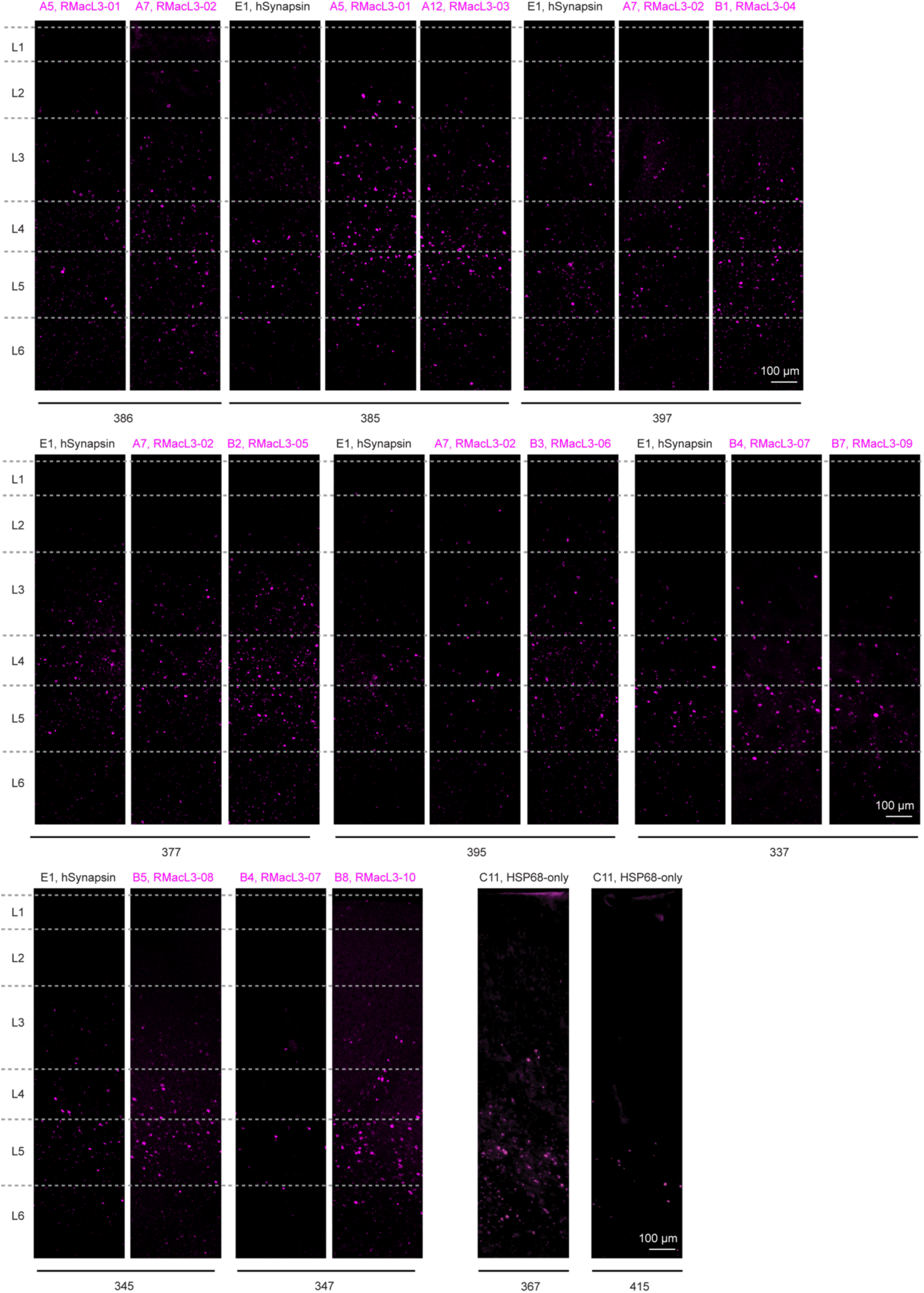
Enhancer screening with FISH, Related to Figure 3. Original FISH images show the enhancer expression in individual sections. The numbers below images indicate brain slice section numbers.

**Figure S4.**
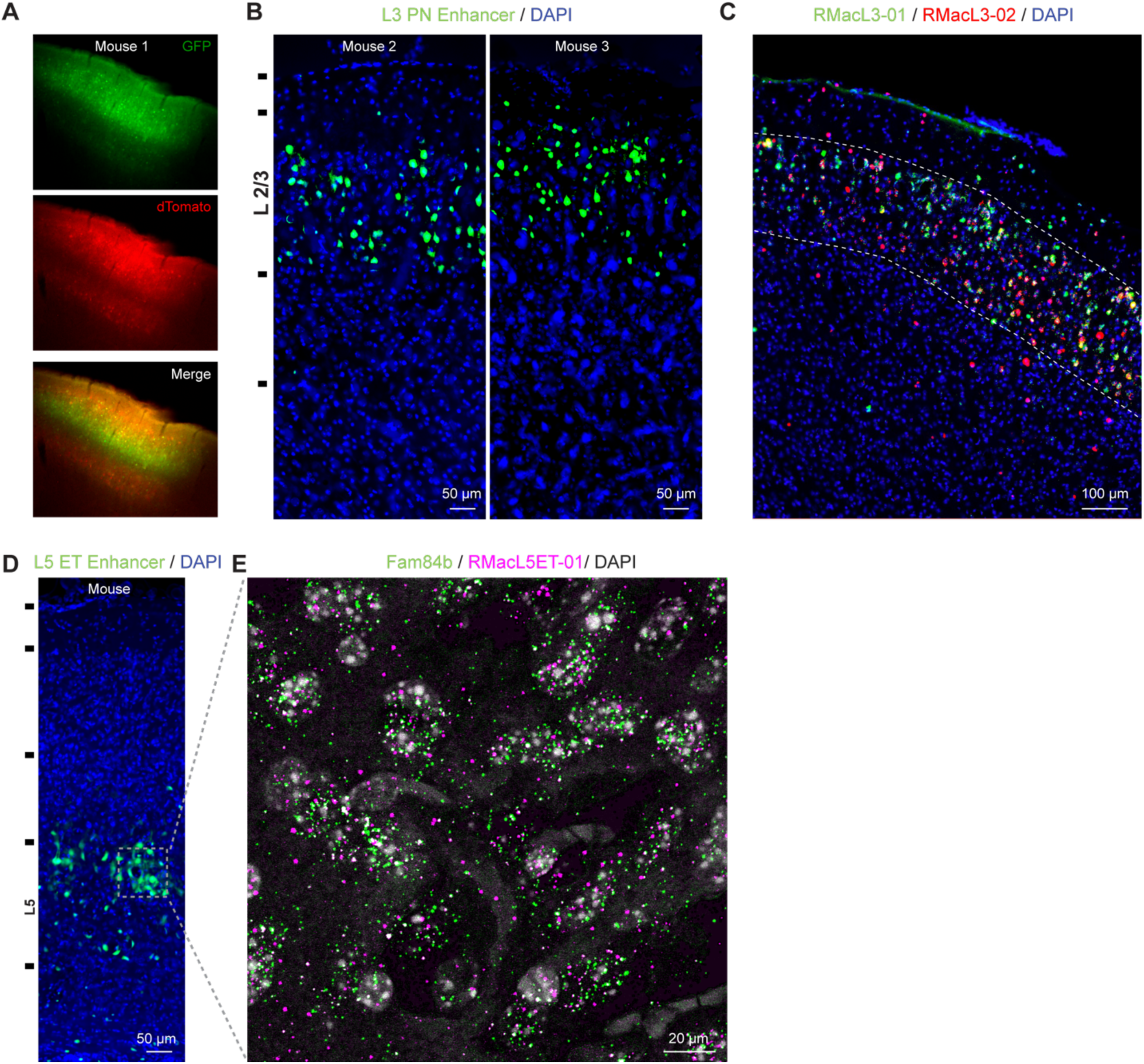
Enhancer expression in rodents, Related to Figure 3. (A) L3PN enhancer-driven GFP expression (library 1) and hSynapsin driven dTomato expression in mouse cortex. (B) L3PN enhancer-driven GFP expression (library 1) in cortical layers of two additional mice. (C) FISH analysis of enhancer expression of RMacL3-01 and −02 showing their expression was enriched in layer 2/3. (D) L5ET enhancer-driven GFP expression (library 1) in mouse cortex. (E) FISH analysis of RMacL5ET-01 and L5ET marker Fam84b shows colocalization.

## STAR*METHODS

### KEY RESOURCES TABLE

| REAGENT or RESOURCE | SOURCE | IDENTIFIER |
| --- | --- | --- |
| Chemicals, Peptides, and Recombinant Proteins |  |  |
| TSA Vivid Fluorophore Kit 520 | Tocris Bioscience | Cat# FP1487A |
| TSA Vivid Fluorophore Kit 570 | Tocris Bioscience | Cat# FP1488A |
| TSA Vivid Fluorophore Kit 650 | Tocris Bioscience | Cat# FP1496A |
| Prolong Gold Antifade Mountant | Life technologies | Cat# P36930 |
| EZ PREP Buffer | Millipore Sigma | Cat# NUC-101 |
| RNase inhibitor | Clontech | Cat# 2313A |
| RNase inhibitor | Roche | Cat# 03335402001 |
| Critical Commercial Assays |  |  |
| Chromium Single Cell 3' GEM, Library & Gel Bead Kit v3.1 | 10X Genomics | Cat# PN-1000121 |
| Chromium Next GEM Single Cell Multiome | 10X Genomics | Cat# PN-1000283 |
| Chromium Single Cell G Chip Kit | 10X Genomics | Cat# PN-1000120 |
| Chromium Single Cell J Chip Kit | 10X Genomics | Cat# PN-1000234 |
| SPRIselect reagent | Beckman Coulter | Cat# B23318 |
| KAPA Library Quantification Kit | KAPA Biosystems | Cat# KK4824 |
| NovaSeq 6000 System | Illumina | Cat# 20013740 |
| NovaSeq S4 Reagent Kit v1.5 (300 cycles) | Illumina | Cat# 20028312 |
| RNAscope Multiplex Fluorescent Detection Kit v2 | Advanced Cell Diagnostics | Cat# 323110 |
| RNAscope Probe: Mmu-POU3F1 | Advanced Cell Diagnostics | Cat# 1005811 |
| RNAscope Probe: Mmu-NR4A2 | Advanced Cell Diagnostics | Cat# 1005861 |
| RNAscope Probe: Mmu-MEIS2 | Advanced Cell Diagnostics | Cat# 1005681 |
| RNAscope Probe: Mmu-CUX2 | Advanced Cell Diagnostics | Cat# 1005671 |
| RNAscope Probe: Mmu-TBX15 | Advanced Cell Diagnostics | Cat# 1005731 |
| RNAscope Probe: Mmu-PCP4 | Advanced Cell Diagnostics | Cat# 100585 |
| RNAscope Probe: Mmu-TYR | Advanced Cell Diagnostics | Cat# 1005719 |
| RNAscope Probe: Mmu-ITGA8 | Advanced Cell Diagnostics | Cat# 1072981 |
| RNAscope Probe: Mmu-NKD1 | Advanced Cell Diagnostics | Cat# 1072991 |
| RNAscope Probe: Mmu-SYT6 | Advanced Cell Diagnostics | Cat# 576481 |
| RNAscope Probe: Mmu-ALPL | Advanced Cell Diagnostics | Cat# 1079991 |
| RNAscope Probe: Mmu-RORB | Advanced Cell Diagnostics | Cat# 876301 |
| RNAscope Probe: Mmu-LAMP5 | Advanced Cell Diagnostics | Cat# 1005831 |
| RNAscope Probe: Mmu-VIP | Advanced Cell Diagnostics | Cat# 870571 |
| RNAscope Probe: Mmu-NDNF | Advanced Cell Diagnostics | Cat# 1075711 |
| RNAscope Probe: Mmu-SST | Advanced Cell Diagnostics | Cat# 461681 |
| RNAscope Probe: Mmu-TH | Advanced Cell Diagnostics | Cat# 1075731 |
| RNAscope Probe: Mmu-PVALB | Advanced Cell Diagnostics | Cat# 461691 |
| Deposited Data |  |  |
| Raw and analyzed data | This paper | Will be available upon publication |
| Experimental Models: Macaque monkeys |  |  |
| Rhesus macaque SM11 | MD Anderson | NA |
| Rhesus macaque SM16 | MD Anderson | NA |
| Rhesus macaque SM20 | UC Davis | NA |
| Rhesus macaque SM19 | UC Davis | NA |
| Rhesus macaque SM45 | UC Davis | NA |
| Rhesus macaque SM18 | Plum | NA |
| Rhesus macaque SM22 | MD Anderson | NA |
| Rhesus macaque SM33 | John Hopkins | NA |
| Rhesus macaque SM35 | John Hopkins | NA |
| Cynomolgus macaque SM38 | World Wide Primates | NA |
| Cynomolgus macaque SM39 | World Wide Primates | NA |
| Cynomolgus macaque SM40 | World Wide Primates | NA |
| Cynomolgus macaque SM41 | World Wide Primates | NA |
| Cynomolgus macaque SM53 | Alpha Genesis | NA |
| Cynomolgus macaque SM63 | World Wide Primates | NA |
| Oligonucleotides |  |  |
| Chromium i7 Multiplex Kit | 10X Genomics | PN-120262 |
| Software and Algorithms |  |  |
| Cell Ranger Analysis Pipeline | 10X Genomics | http://10xgenomics.com/ |
| Seurat |  | https://satijalab.org/seurat/ |
| Scanpy |  | https://scanpy.readthedocs.io/en/stable/ |
| ImageJ | Schneider et al., 2012 | https://imagej.nih.gov/ij/ |
| CellProfiler 3.1.9 | Carpenter et al., 2006 | https://github.com/CellProfiler |
| Analysis pipeline | This paper |  |

### RESOURCE AVAILABILITY

Reagents and resources should be directed to the lead contact.

### EXPERIMENTAL MODEL AND SUBJECT DETAILS

#### NHPs

All animal procedures were in accordance with the National Institutes of Health Guide for the Care and Use of Laboratory Animals and approved by the University of Pittsburgh’s Institutional Animal Care and Use Committee (IACUC) (Protocol ID, 19024431 and 21058910). Rhesus and Cynomolgus Monkeys were single- or pair-housed with a 12h-12h light-dark cycle. Monkey SM11 was a 4-year-old male (5.6 kg). Monkey SM16 was a 2.5-year-old male (3.9 kg). Monkey SM20 was a 3-year-old female (4.5 kg). Monkey SM19 was a 3-year-old male (4.9 kg). Monkey SM45 was a 3-year-old male (4.5 kg). Monkey SM33 was a 20-year-old male (3.26 kg). Monkey SM35 was a 2-year-old female (4.03 kg). Monkey SM41 was a 9-year-old male (6.86 kg). Monkey SM53 was a 6-year-old male (6.13 kg). Monkey SM38 was a 10-year-old male (10.9 kg). Monkey SM39 was a 12-year-old male (9.26 kg). Monkey SM40 was an 11-year-old male (8.26 kg). Monkey SM63 was a 5-year-old male (6.73 kg). For FISH: Monkey SM18 was a 12-year-old male (8.5 kg); Monkey SM22 was a 3-year-old male (5.2 kg).

### METHOD DETAILS

#### Brain surgery for single cell experiments

We performed brain surgery as previously described ^52^. Briefly, animals (monkeys SM11, SM16, SM19, SM20, SM45) were anesthetized with ketamine (15 mg/kg IM) within a cage and transported to a surgery suite. The animals were further kept anesthetized with isoflurane and head-fixed in a stereotaxic instrument (Kopf Instruments). Vitals, including body temperature and respirations, were under constant monitoring. All surgical tools were sterilized and wiped with RNase decontaminant (Apex, Cat# 10-228). To maximize the viability of the cells, we removed the skull first before perfusing the monkeys with approximately 4 liters of oxygenated ice-cold artificial cerebrospinal fluid. We then opened the dura and removed the brain. We cut the prefrontal cortex under a dissection microscope for nuclei isolation. Monkeys SM18 and SM22, for FISH, were perfused first with phosphate-buffered saline (PBS, Fisher Scientific, Cat# BP243820) to flush out blood and then with 4% paraformaldehyde (PFA, Sigma-Aldrich, Cat# P6148) supplemented with 10% sucrose (Sigma-Aldrich, Cat# S8501) to fix the tissue. The brain was post-fixed with 4% PFA and cryopreserved with a gradient of sucrose (10%, 20%, 30%) in PBS.

#### Brain surgery for viral injections

We scanned monkeys with T1 weighted scan using a Siemens MAGNETOM 7T Plus at least one week before surgery. These scans (between 3 and 5 individual scans per animal) were later re-oriented into the coronal plane and averaged before loading into Brainsight®. This approach allows for accurate surgical targeting in the absence of a stereotaxic MRI. When the averaged and oriented scan was loaded into Brainsight, the program allowed for 3D image reconstruction based on the MRI. Using landmarks on the skull and a 3D brain model, our intended targets were located and programmed prior to surgery.

On the day of surgery, we placed monkeys into a stereotaxic frame to mimic the coronal positioning of the MRI scan. A sagittal opening was made in the skin and fascia using a cauterizing tool. We opened the temporalis muscle’s internal capsule to enable muscle retraction for adequate access to the target sites. A laser pointer was attached to the surgical robot, and two camera capture points were projected on the exposed skull. This registration was compared to the 3D rendering created from the MRI file prior to surgery. The program could then overlay the 3D rendering and planned targets to the monkey’s exposed skull for targeting. Once registration was complete, we removed the laser, drilled burr holes, and infused viruses at a slow controlled rate with a Harvard Apparatus Pump 11 elite Nanomite. Each track generally consisted of multiple injection sites at different dorsal/ventral locations until the most dorsal injection site was reached. At the end of the track, the needle stayed in place for 5-10 minutes to allow the virus to diffuse into the target tissue before being removed from the tissue. After the end of all injections, we closed soft tissue in anatomical layers from muscle capsule to skin. All procedures were conducted using aseptic techniques in a dedicated operating suite. At least 12 hours before surgery, subjects were treated with antibiotics (ceftriaxone; 50mg/kg at 350mg/mL, IM) and steroidal anti-inflammatory (dexamethasone; 0.5mg/kg at 4mg, PO) to minimize the risk of post-operative infection and inflammation. To minimize nausea, we gave Cerenia (1mg/kg at 10mg/mL, SQ) the night before and the night of the surgical procedure. Antibiotic and steroid treatment was continued for 3-7 days post-surgery along with analgesics such as Meloxicam (0.2mg/kg at 1.5mg, PO) and Buprenorphine (0.02mg/kg at 0.4mg/mL, IM). For the one-at-a-time validation experiments, the injection details are found in Table S2

#### Optogenetics

On the day of surgery, monkeys were placed into a stereotaxic frame, and the opening was the same as that for injection surgeries. However, once the skull was cleaned, we opened a rectangular window over the cortex, which had been injected with the virus in the previous procedure. Using surgical blade #11, an incision was made in the dura, exposing the cortex of interest. To stabilize cortical pulsations, 3% agarose (Invitrogen) solution was coated over the exposed tissue during data acquisition. We collected optogenetic stimulation and recording using a Plexon custom S-Probe, 16 recording channels with four light channels. The probe was connected to a Saphire 488-500 LPX LDRH Laser System with an LC adaptor. The laser was paired with an adjustable and focusable laser beam coupler and a single-mode fiber cable. Stimulations were fed into the laser coupler from a A-M Systems Isolated Pulse Stimulator (Model 2100). A record of the stimulations and recordings was saved using the Spike2 Program and later analyzed with MATLAB. Once all the data were collected, we closed the soft tissue and prepared the animal for perfusion. All procedures were conducted using aseptic techniques in a dedicated operating suite. Isoflurane sedation was kept at a minimum during the procedure, with additional doses of ketamine (15mg/kg at 100mg/mL) provided every 6 hours.

#### Single nucleus RNA sequencing

We ran 10x Chromium Single Cell 3’ Reagent kits, v3.1 Chemistry (10x Genomics, Cat# PN-1000121) for monkeys SM11 and SM16. We isolated nuclei as previously described. ^52^ We followed the standard 10x protocol for v3.1 chemistry for library generation. Briefly, we reverse transcribed mRNAs within Gel beads-in-emulsion (GEMs) after running through a 10x Genomics Chromium controller. We broke the emulsion with a recovery agent (10x Genomics, Cat# 220016) and purified cDNAs with Dynabeads (10x Genomics, Cat# 2000048). Then, we amplified cDNAs and purified them with SPRIselect reagent (Beckman Coulter, Cat# B23318). After analyzing the cDNA quality using an Agilent Bioanalyzer 2100, we prepared libraries following fragmentation, end repair, A-tailing, adaptor ligation, and sample index PCR. We quantified the libraries by qPCR using a KAPA Library Quantification Kit (KAPA Biosystems, Cat# KK4824). We pooled together libraries from individual monkeys and loaded them onto NovaSeq S4 Flow Cell Chip. We sequenced samples to the depth of 200,000 reads per nucleus.

#### Single nucleus RNA and ATAC multiomics sequencing

We used the 10x Chromium Single Cell Multiome Library & Gel Bead Kit (10X Genomics, Cat# PN-1000283) for monkeys SM19, SM20, and SM45. Briefly, we broke the tissue into small pieces with a wide-bore pipette tip followed by a regular-bore pipette tip, then filtered through a 30 μm MACS SmartStrainer. After spin down, we lysed the cells with 0.1X lysis buffer (10mM Tris-Hcl, 10mM NaCl, 3mM MgCl_2_, 0.1% Tween-20, 0.1% NP-40, 0.01% Digitonin, 1% BSA, 1mM DTT, 1U/ul RNase inhibitor). After washing, we resuspended the nuclei with diluted nuclei buffer (10x Genomics, PN-2000153). We adjusted the nuclei concentration to 2,900-7,260 nuclei/μl, based on the targeted nuclei recovery of 9,000. We transposed the nuclei and ran through the 10x chromium controller, followed by reverse transcription. We generated the snATAC-Seq and snRNA-Seq libraries according to the standard protocol. We pooled together libraries and loaded them onto a NovaSeq 6000 S4 Flow Cell. We sequenced each sample from Monkeys SM19, SM20, and SM45 to the depth of 150,000 x 150 bp paired-end reads per nuclei.

#### snRNA-Seq analysis

After sequencing, we converted the BCL reads to fastq files and aligned the reads to a custom transcriptome reference. We integrated the monkey dataset using a standard Seurat v3 pipeline. Briefly, we first removed ambient RNA, ribosomal genes, and doublets. Then we performed standard log-normalization and a variance stabilizing transformation and identified variable features individually for each monkey’s dataset using Seurat’s FindVariableFeatures function. Next, we identified anchors using the FindIntegrationAnchors function with default parameters and passed these anchors to the IntegrateData function. This returned a Seurat object with an integrated expression matrix for all nuclei. We scaled the integrated data with the ScaleData function, ran PCA using the RunPCA function, and visualized the results with UMAP. We used Louvain clustering and chose a resolution that reflected the major cell classes of the PFC, including excitatory neurons, inhibitory neurons, and astrocytes. We calculated the differentially expressed genes for each cell class with the FindMarkers function and, based on the marker genes, we annotated the major cell classes.

Because the variation with excitatory neurons was masked when co-clustering with other cell types, we isolated the excitatory neuron clusters that express excitatory neuron marker genes such as SLC17A7 and TBR1. We re-calculated the principal components (PCs) and performed UMAP dimension reduction and chose a resolution that separated clusters that were distinct in UMAP space. We annotated these clusters based on their maker genes and mapping to laminar locations. To analyze the inhibitory neuron populations, we isolated clusters based on inhibitory neuron markers *GAD1* and *GAD2*. Similarly, we re-calculated PCs and performed UMAP dimensionality reduction on the first 20 PCs. We used Louvain clustering and annotated the resulting clusters based on known interneuron markers.

We calculated the cosine similarity for excitatory and inhibitory neurons within and between clusters based on PCA space. We used a permutation test on the cosine similarity between pairs of clusters. We randomly shuffled the nuclei to mask the nuclei identity and recalculated cosine similarity. We repeated these 10,001 times and used within group and between group variance ratio to determine a *P* value. To compare the cell types between monkey and human excitatory neurons, we transferred the labeling of human PFC, MTG, and M1 subclass and cell type to our reference monkey dataset with Seurat *TransferData* function. We normalized the cell number for each human subclass/cell type and plotted a heatmap to show their correspondence relationship.

#### snATAC-Seq alignment and cell quality control (QC) filtering

We aligned snATAC-Seq reads from all samples to the rheMac10 genome using the fast snATAC-Seq mapper chromap.^53,54^ We created arrow files from the fragment files for downstream analyses with the ArchR package.^55^ We followed the previous approach to map the higher quality GRCh38.p13 human RefSeq gene annotations onto the rheMac10 genome with the liftOff tool and used this to create a custom rheMac10 ArchR gene and genome annotation.^52,55,56^ Using this liftOff annotation, we were able to compute ArchR gene activity scores around the orthologous rhesus macaque regions of human genes, enabling us to obtain scores for substantially more genes and complete gene bodies than we would have been able to obtain using base rheMac10 gene annotations. We excluded genes, exons, and transcription start sites that are within 1Mb from chromosome ends which creates errors during per-cell gene score computation. Within ArchR, we performed two rounds of low-quality nuclei removal. This 2-step filtering process excludes 6-12% of unfiltered nuclei and removes nuclei that tend to form clusters entirely made of low-QC cells or single nuclei that do not cluster with other high-quality nuclei. The first round used cut-offs to exclude nuclei with a high probability of being doublets (cutEnrich = 0.5, cutScore = -log10(.05), filterRatio = 1) and low number of unique per cell fragments (nFrags < 10^3.5). The second round used the MASS R-package negative-binomial generalized linear model to learn the relationship between per-cell metrics TSSenrichment and PromoterRatio and DoubletEnrichment with the number of unique fragments for each sample, glm.nb(nFrags ∼ TSSEnrichment + PromoterRatio + DoubletEnrichment).^57^ We calculated the standardized residual from the curve fit and excluded nuclei with residuals more than two standard deviations away from the fit.

#### Single nuclei ATAC clustering, cell annotation, and differential analyses

We performed iterative latent-semantic index clustering with 4 iterations and 30,000-230,000 variable features, followed by Harmony batch correction,^58^ UMAP visualization, and Louvain clustering, where we used default settings for all methods. We labeled snATAC-Seq clusters by first identifying glial and neuronal clusters by marker gene activity scores (*RBFOX3*, *GAD1*, *GAD2*, *AQP4*, *CX3CR1*, *PDGFRA*, *MOG*, etc.). We subset glial and neuronal clusters and re-performed iterative LSI embedding and clustering as above. To annotate the snATAC-Seq nuclei, we applied Seurat’s canonical correlation analysis (CCA) on ArchR gene activity scores with the corresponding RNA profiles from the same animal and brain region, which was confirmed with the barcode link associated with the multiomics kit. We created pseudo-bulk profiles across the annotated cell types and replicates with ArchR function addGroupCoverage(minReplicates = 5, maxReplicates = 24, minCells = 40, maxCells = 1000), called peaks across replicates with macs2,^59^ identified reproducible peaks between replicates for each cell type, created a consensus peak set of 501bp fixed-width, summit-centered peaks across cell types, and created a peak by cell, “PeakMatrix”, through ArchR.

#### Machine models of cell type specificity

We further split peaks into the test set, validation set, and training set. We trained support vector machine (SVM) models with lsgkm gkmpredict function with the parameters -t 4 -l 7 -k 6 -d 1 -M 50 -H 50 -m 40000 -s -T 4 previously found to be robust to train accurate models.^26,60^ We performed a grid search for the regularization -c and weight -w parameters to train SVM models and selected the models with the best F_half score evaluated on the validation set. We excluded models that did not have area under the receiver operator characteristic (auROC) curve > .65 and accuracy > .65 on the validation set. We trained convolutional neural network (CNN) models with the 5-convolutional layer architecture and one-cycle policy training regime previously described.^61^ We used a batch size of 64 sequences and 30 epochs. We excluded models that did not have area under the auROC curve > .65 and accuracy > .65 on the validation set.

#### Prioritization of cell type specific enhancers

We scored differential peaks for each cell type with the collection of best SVMs and CNNs successfully trained for each comparison and averaged scores from both SVM and CNN for the same comparison and across comparisons to rank the degree of cell type specificity. We screened out peaks that were predicted to have off-target effects in any other cell type by removing any peak with negative predictions across comparisons. In addition, for each candidate enhancer, we identified whether the accessibility of these enhancers correlated with increased gene expression of nearby genes. Using the subset of nuclei with same-cell ATAC and RNA profiles, we used the ArchR Peak2GeneLinkage() function to identify peaks statistically co-accessible with measured gene expression. Furthermore, we added further priority to peaks where the associated gene was also separately identified as a differentially expressed marker gene. We applied the geometric mean to the Peak2Gene correlation with gene expression, MarkerGene log2-fold change, cell type specific motif Z-score, and the average ML score to create the composite score of whether the candidate peak contains cell type specific DNA sequences based on ML and motif analyses likely drives expression of cell type specific genes by co-accessibility and differentially expressed gene analyses. Finally, we inspected genome track plots of the open chromatin around the candidate enhancer and gene expression of nearby genes to select the top 12 candidates to test for cell type specificity in the rhesus macaque brain.

#### Plasmid Cloning and AAV purification

To test the effects of these putative cell type specific enhancers, we generated plasmids with each enhancer upstream of the minimal promoter HSP68 to drive the expression of GFP. A control plasmid was also generated in which the human synapsin promoter drove the expression of dTomato. For all plasmids, we put a 500bp barcode downstream of the transgene to allow for FISH analysis of activity. These inserts were synthesized and cloned into backbone pEMS2115 (Addgene Plasmid #49140) with restriction enzymes EcoRI and Not1. Plasmids were packed into AAV9 2YF, PHP.eB, and X1.1 serotypes for viral purification (Vectorbuilder).

#### FISH stain and imaging

We removed the excess solution from the brain surface, air-dried the brain for 15 min, embedded brains in optimal cutting temperature compound (OCT), and stored them at −80°C until sectioning. We collected 15 μm free-floating sections from monkeys SM18 and SM22 and mounted them on 2×3” pre-coated slides. We preserved the mounted sections in a −80°C freezer. We performed FISH with Multiplex Fluorescent Detection Reagents v2 (ACD, Cat# 323110) according to the manufacturer’s protocol with slight modifications for monkey brain tissue. We retrieved slides from −80 °C freezer and equilibrated the slides to room temperature for 30 min. We washed the slides briefly with water and fixed the tissue in 4% PFA buffer for 30 min. The sections were then dehydrated in 50%, 70%, and 100% ethanol and baked at 60 °C for 20 min. We incubated the brain sections with hydrogen peroxide for 10 min to quench endogenous horseradish peroxidase (ACD, Cat# 322335) and then performed target retrieval with RNAscope Target Retrieval Reagents (ACD, Cat# 322000) for 8 min at 99°C. We dehydrated the slides in 100% alcohol for 3 min and then baked them at 60°C for 10 min. We incubated the slides with protease III (ACD, Cat# 322337) for 30 min at 40°C to increase probe penetration and then hybridized them with probes for 2 hr. After signal amplification with AMP 1, 2, and 3, we conjugated probes with different HRP channels and fluorophores, including Opal 520 (PerkinElmer, Cat# FP1487A), Opal 570 (PerkinElmer, Cat# FP1488A), and Opal 650 (PerkinElmer, Cat# FP1496A). We counterstained sections with DAPI and mounted them with Prolong Gold Antifade Mountant (Life technologies, Cat# P36930). We scanned sections using a Nikon Eclipse T*i*2 under 20x objective. We used ImageJ and Adobe Photoshop to adjust brightness and overlay images.

### QUANTIFICATION AND STATISTICAL ANALYSIS

#### Enhancer distribution across layers

To assess the enhancer expression across layers, we used Cellpose to detect the cells and quantify their distribution.^62^ Briefly, we first adjusted the orientation of the images to be vertical and of equal width. Then we ran Cellpose to detect the cells using the default parameters. Cellpose output the positions, sizes, and signal intensities of cells. We set a consistent threshold for size and signal intensity to remove noise signals. We divided the layers into 50 parts and counted the number of cells in each segment. We normalized the cell number to the maximum number for each image and plotted their layer distribution. The layer boundaries are judged based on the original DAPI images.

#### Enhancer cell type specificity/efficiency by colocalization with marker genes

To determine the specificity and efficiency of enhancer-driven expression, we performed FISH against enhancer-specific barcodes and marker genes. Using CellProfiler cell image analysis software (cellprofiler.org), intensities and locations for each RNA signal were defined for each cell. We performed cell counts for colocalization of FISH signal for the RMacL3-01 barcode and *CUX2*. Specificity was calculated as the percentage of enhancer barcode-expressing cells that also co-expressed the *CUX2*. Efficiency was calculated as the percentage of *CUX2*-expressing cells that also co-expressed the enhancer barcode. Results are reported as mean ± SD (n=4).

#### Brain-wide FISH mapping

Our image reconstruction approach uses a generative probabilistic model incorporating diffeomorphic spatial warping and contrast adjustments. Mapping to a common coordinate system is achieved through maximum a posteriori estimation, enabling precise reconstruction of both 3D and 2D datasets. To minimize tissue distortion, we employed the Tape-Transfer cryosectioning method,^63,64^ which preserves the tissue’s original spatial orientation with 99% distortion-free accuracy.^63^ This framework supports data from MRI, various staining methods, and partial tissue regions. By integrating different types of histology and FISH data, our workflow provides a unified coordinate framework for the quantification of ground-truth distribution across datasets for brain mapping. We first preprocessed histology sections (downsample) to match MRI resolution using scattering operators. This approach preserves critical texture information necessary for accurate multimodal alignment. We then adopted a Rigid Alignment from each image acquisitions to minimize a multimodal cost function, which accounts for contrast transformations and the likelihood of pixel artifacts. To address signal inhomogeneity and large inter-modality differences, we optimized parameters locally in small regional blocks. The optimization was carried out iteratively with a multi-resolution approach to avoid local minima. *Initial 3D Reconstruction*: We reconstructed the 3D structure of histology sections and regional tissues by rigidly aligning them with neighboring sections. This alignment was adjusted to account for any anomalous pixels, ensuring accurate reconstruction. *MRI-guided Registration*: Diffeomorphic mappings between MRI and histology sections were created using a sequence of spatial warps based on the Large Deformation Diffeomorphic Metric Mapping (LDDMM) model. ^65^ *Multimodal Registration*: FISH data was then aligned to the transformed histology sections using rigid alignment. Partial samples could also be aligned manually with a rigid initialization framework if needed. All transformations between spatial domains were saved in a standardized format. Throughout the process, we measured uncertainties by positioning corresponding points, curves, and surfaces and by evaluating the true distances between them for further analysis.

